# Identification of Glyoxalase A in Group B *Streptococcus* and its contribution to methylglyoxal tolerance and virulence

**DOI:** 10.1101/2024.07.30.605887

**Authors:** Madeline S. Akbari, Luke R. Joyce, Brady L. Spencer, Amanda Brady, Kevin S. McIver, Kelly S. Doran

## Abstract

Group B *Streptococcus* (GBS) is a Gram-positive pathobiont that commonly colonizes the gastrointestinal and lower female genital tracts but can cause sepsis and pneumonia in newborns and is a leading cause of neonatal meningitis. Despite the resulting disease severity, the pathogenesis of GBS is not completely understood, especially during the early phases of infection. To investigate GBS factors necessary for blood stream survival, we performed a transposon (Tn) mutant screen in our bacteremia infection model using a GBS *mariner* transposon mutant library previously developed by our group. We identified significantly underrepresented mutations in 623 genes that contribute to survival in the blood, including those encoding known virulence factors such as capsule, the β-hemolysin, and inorganic metal ion transport systems. Most of the underrepresented genes have not been previously characterized or studied in GBS, including *gloA* and *gloB,* which are homologs for genes involved in methylglyoxal (MG) detoxification. MG is a byproduct of glycolysis and a highly reactive toxic aldehyde that is elevated in immune cells during infection. Here, we observed MG sensitivity across multiple GBS isolates and confirm that *gloA* contributes to MG tolerance and invasive GBS infection. We show specifically that *gloA* contributes to GBS survival in the presence of neutrophils and depleting neutrophils in mice abrogates the decreased survival and infection of the *gloA* mutant. The requirement of the glyoxalase pathway during GBS infection suggests that MG detoxification is important for bacterial survival during host-pathogen interactions.

## Introduction

*Streptococcus agalactiae* (group B *Streptococcus*, GBS) is an opportunistic pathogen that commonly resides in the gastrointestinal and lower female genital tracts but can cause infection in newborns and is also increasingly associated with non-pregnant individuals, especially older adults and patients with diabetes (*1–3*). GBS asymptomatically colonizes the vaginal tract in up to 30% of people but can instigate complications during pregnancy and birth, such as preterm labor, and serious infections in newborns, such as sepsis, pneumonia, and meningitis (*1, 4–6*). Research into GBS intrauterine infection during pregnancy thus far indicates that GBS-activated inflammatory pathways ultimately result in preterm births (*7*). If GBS is vertically transferred to the neonate, the resulting infection is categorized as either early-onset disease (EOD, 0-6 days of life) or late-onset disease (LOD, 7-90 days of life) depending on the timing of symptom presentation (*8*). Neonatal meningitis caused by both EOD and LOD GBS disease requires a sustained level of bacteremia prior to the penetration into the brain and, even after treatment, frequently results in long-lasting neurological effects and long-term morbidity (*4, 5*). Although intrapartum antibiotic prophylaxis is administered to colonized pregnant women to prevent the detrimental effects of GBS infection, GBS isolates are increasing in resistance to second-line antibiotics over time (*9*) and intrapartum antibiotic prophylaxis is not effective in preventing LOD. Therefore, studying GBS pathogenesis of meningitis, including bacteremia, is important for the development of novel treatments and therapeutics to prevent GBS infection and reduce morbidity and mortality.

Previous work has determined the GBS transcriptome as well as genes necessary for survival in human blood *in vitro* and for survival of the murine female reproductive tract (*10–13*). These datasets as well as other studies have shown that GBS possesses an arsenal of virulence factors that directly contribute to pathogenesis such as β-hemolysin/cytolysin, superoxide dismutase, capsule, adherence proteins, and metal transport systems (*10, 14, 15*). β-hemolysin/cytolysin (βH/C) and capsular polysaccharide are the most well studied factors associated with GBS pathogenesis and are regulated by the well-known two-component system, CovR/S (*15*). βH/C is an ornithine rhamnolipid pigment synthesized by the *cyl* operon and has both hemolytic and cytolytic capabilities against a variety of host cells including red blood cells, neutrophils, macrophages, and epithelial cells (*16–19*). As a result, βH/C has been shown to contribute to GBS blood, lung, and brain infection (*17, 20*). Capsular polysaccharide is surface-associated and made up of different arrangements of monosaccharides that form capsular repeat units (*21, 22*). There are 10 known GBS capsular serotypes with serotype III being highly associated with neonatal infections, such as meningitis, and which is overrepresented in invasive isolates worldwide (*9, 23, 24*). Group B streptococcal capsular polysaccharide was first studied over 40 years ago and has been shown to help GBS evade host immune defenses by mimicking host antigens and blocking complement-mediated opsonophagocytic killing as well as to facilitate GBS biofilm formation (*22, 25–27*). Despite these studies, the contribution of GBS metabolism to colonization and infection *in vivo* has been a neglected area of study in the field (*10, 15*).

Here we performed a transposon-mutant screen (Tn-sequencing) using a murine bacteremia model to discover additional genes necessary for GBS fitness in murine blood *in vivo*. GBS survival within the blood is an essential prerequisite to penetrate the blood brain barrier and subsequence development of meningitis. Tn-sequencing allows for the identification of genes that may be continuously expressed but are essential in certain environments. Here, we identify that the glyoxalase pathway is required for GBS bloodstream survival. The glyoxalase pathway consists of two genes, *gloA* and *gloB,* and is involved in methylglyoxal detoxification (*28, 29*). Methylglyoxal (MG) is toxic byproduct of normal cell metabolism (*30*), and we confirm that the first enzyme in the pathway, encoded by *gloA,* contributes to GBS MG detoxification and invasive infection. Furthermore, we found that *gloA* is necessary for GBS neutrophil survival and depleting neutrophils rescues the *gloA* mutant *in vivo*.

## Results

### Genome-wide analysis of GBS factors involved in bloodstream survival

To identify genes necessary for GBS survival in murine blood, we utilized a bacteremia model of infection with our previously described Tn mutant library in the CJB111 strain, a GBS isolate from a case of neonatal bacteremia without focus (*17, 31–33*). Briefly, mice were intravenously infected with the Tn mutant library and the infection was allowed to progress up to 27 hours. The input Tn mutant library and libraries recovered from the blood were processed as described in Materials and Methods. To identify transposon insertion sites, sequenced reads were mapped to the GBS CJB111 genome, which identified 623 genes as significantly underrepresented (p_adj_ < 0.05, log_2_FC < –1) and 95 genes as significantly overrepresented (p_adj_ < 0.05, log_2_FC > 1) in the blood compared to the input library (**Fig. 1A**) (**Table S1**). The significant gene hits were equally distributed across the genome. Significant genes were then assigned clusters of orthologous groups of proteins (COGs). The number of significant gene hits in each COG were normalized to the total number of genes in each COG revealing sRNA, amino acid transport and metabolism, and inorganic ion transport and metabolism as the COGs containing the most underrepresented genes during GBS survival in the blood (**Fig. 1B**). We detected many classes of GBS virulence factors and genes known to contribute to GBS infection as significantly underrepresented (**Table 1).** Some of these genes are involved in hemolytic pigment biosynthesis, capsule biosynthesis, two-component regulatory systems, metal transport, glutamine transport, and purine metabolism. When we investigated other underrepresented genes that have not been previously characterized in GBS, we found homologous genes to glyoxalase A and B of the glyoxalase pathway to both be significantly underrepresented with fold changes of –18.38 and –25.63, respectively (**Table 1**). The glyoxalase pathway is a ubiquitous two-step process found across all kingdoms of life and is the primary mechanism of methylglyoxal (MG) breakdown (**Fig. 1C**) (*28*). A highly reactive carbonyl byproduct of normal cell metabolism, most MG is primarily generated from glycolysis, but can also be produced from other metabolic pathways such as lipid and protein metabolism (*34*).

**Figure 1.**
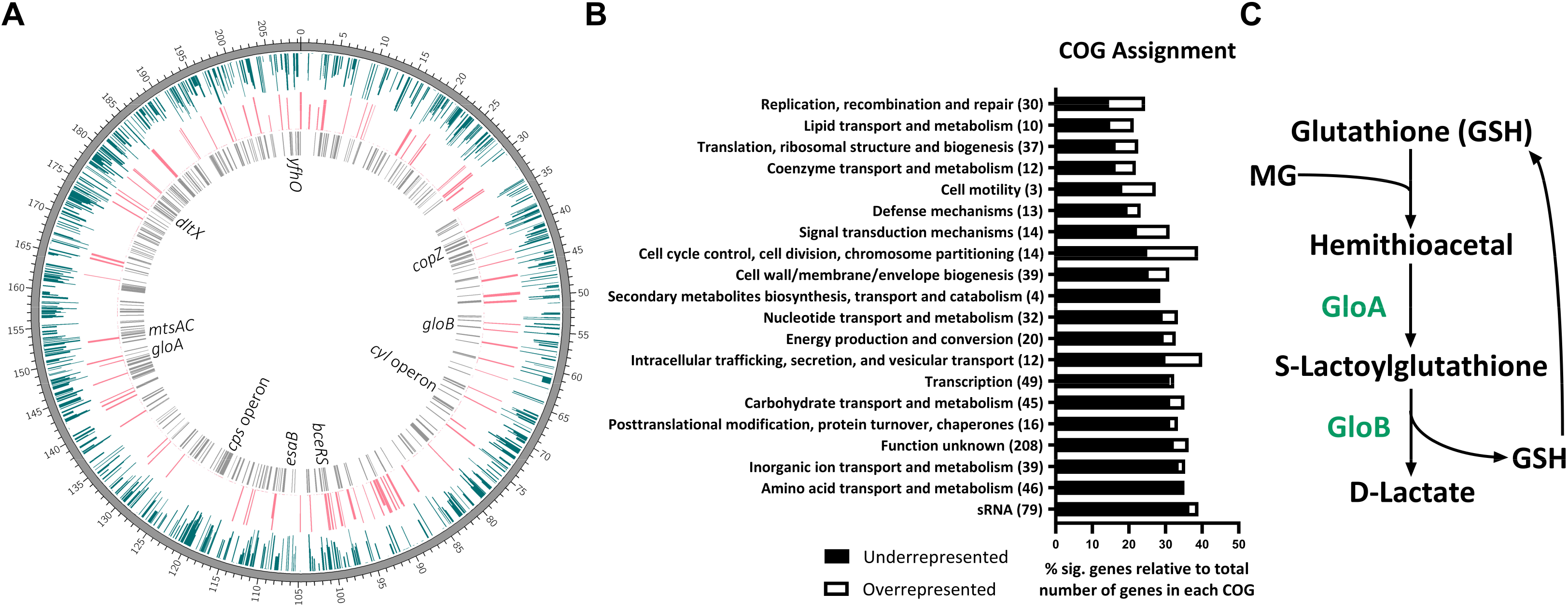
*In vivo* transposon mutant sequencing of GBS survival in the blood. (A) CIRCOS atlas representation of the GBS CJB111 genome is shown with base pair ruler on the outer ring. The inner ring in blue shows log_2_FC (0 to –5/max) of significantly underrepresented genes (P_adj_ < 0.05). The next inner ring in pink shows log_2_FC (0 to 5/max) of significantly overrepresented genes (P_adj_ < 0.05). The most inner ring in grey denotes genes with P_adj_ < 0.001. Underrepresented genes or operons of interest are labeled in the center (also listed in Table 1). (B) Clusters of orthologous genes (COG) assignments for significant gene hits normalized to the total number of GBS genes in each COG. The total number of significant genes in each COG are in parentheses. (C) Diagram of the glyoxalase pathway for methylglyoxal (MG) breakdown. Significance determined by (A&B) TRANSIT analysis and Trimmed Total Reads (TTR) normalization with *P_adj_* < 0.05 and Log_2_ Fold Change < –2 or > 2.

**Table 1.**
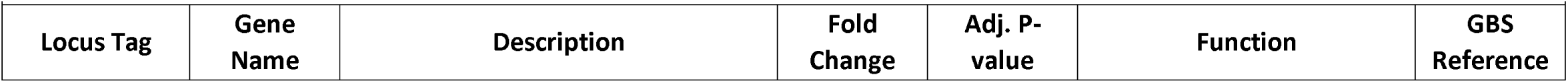
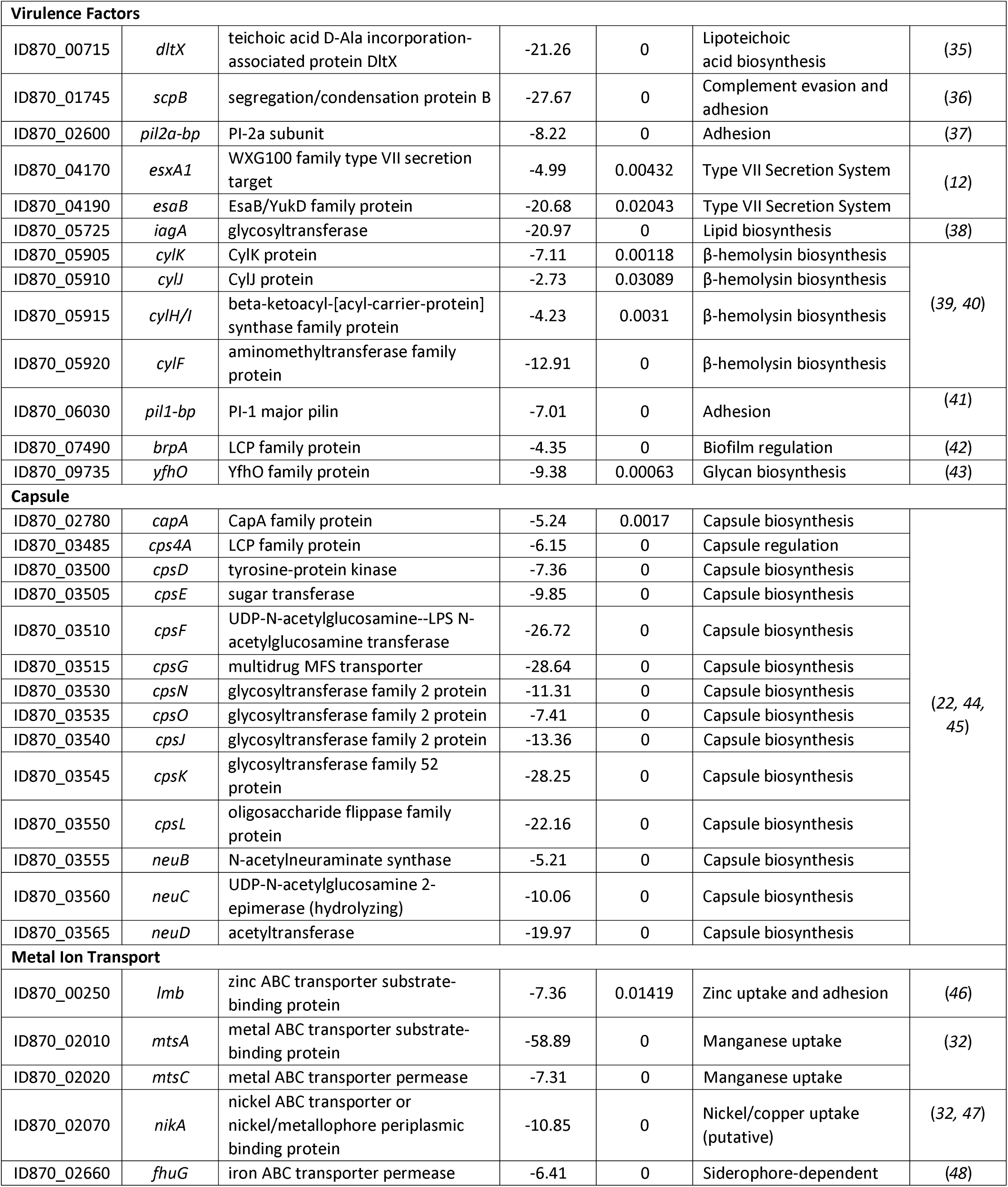
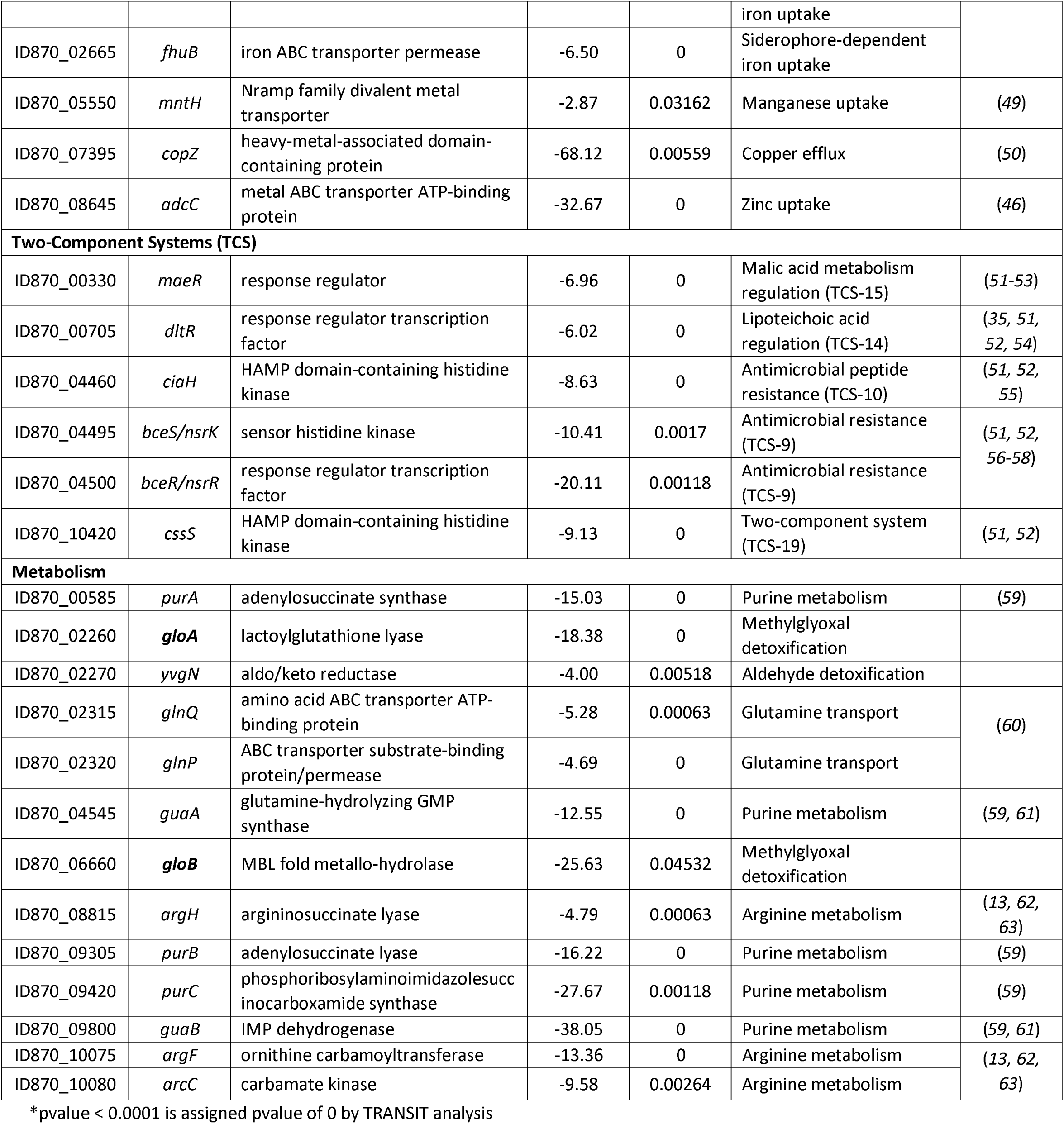
Important GBS virulence factors contribute to survival in blood.

### Methylglyoxal tolerance differs across GBS strains

GBS contains glyoxalase A and B homologs, also known as lactoylglutathione lyase (ID870_02260) and hydroxyacylglutathione hydrolase (ID870_06660) respectively. These are hypothesized to be involved in MG detoxification and therefore, tolerance. To begin to characterize this pathway in GBS, we grew several clinical GBS isolates, representing various capsule serotypes, in the presence of 1.0 mM MG in a modified chemically defined media (mCDM) (*64*) (**Fig. 2**). These concentrations of MG resulted in an observed increase in lag phase without a decrease in CFU, suggesting MG has bacteriostatic properties (**Fig. S1A**). CJB111 exhibited the highest sensitivity to 1.0 mM MG with the largest increase in lag phase quantified by the change in time to max OD_600_ at 8.00 hrs. All of the strains tested had significantly decreased time to max OD_600_ when compared to CJB111, and COH1 exhibited the lowest sensitivity with the smallest time to max OD_600_ at 1.83 hrs. Overall, different GBS isolates had varying degrees of resistance to MG, but resistance was not correlated with serotype or growth rate differences (**Fig. S1B**). To explore the strain differences in MG tolerance further, we selected three representative serotype strains commonly used by our group and others working on GBS pathogenesis with low to high resistance: CJB111 (V), A909 (Ia), and COH1 (III). First, we compared GloA amino acid sequences between these three strains and previously characterized GloA from *Streptococcus pyogenes* and *Escherichia coli* (**Fig. S2A**) (*29, 65–67*). We chose these strains since the GAS glyoxalase pathway has been investigated previously and it is within the same genus and the *E. coli* GloA has a solved crystal structure which is important for protein modeling. The CJB111 GloA is 63.5% identical to GAS GloA and 40.5% identical to *E. coli* GloA. Interestingly, the GloA from CJB111 and A909 are 100% identical while the COH1 GloA is only 99% identical due to a single amino acid change of an alanine to a serine (A45S) in a non-conserved region. To determine how common this variant was in GBS, we generated a phylogenetic tree using BlastP and FigTree to compare 57 GBS genomes and found 12 out of 57 GloA proteins (21%) have the A45S change with another 5 having a different A45 variant (**Fig. S2B**). Proteins with the A45S variant also clustered together in the tree suggesting a common ancestral strain, however, only COH1 from the strains tested in Fig. 2 had this variant. To assess tertiary structure, a predicted protein model for GBS GloA was generated using AlphaFold2 and had extremely high confidence for most residues (**Fig. S2C&D**). The predicted structure was compared to the solved *E. coli* GloA structure (PDB 19FZ) and found to have highly similar topology and conserved metal binding residues (**Fig. S2C**). GloA was also modeled in its active form as a dimer to show predicted active sites (**Fig. S2E Left**). The A45S variant from COH1 GloA was included in the dimer and modeled to be next to the predicted active site (**Fig. S2E Right**). Lastly, using our selected representative strains, we investigated baseline transcription regulation of the glyoxalase pathway by comparing mid-log transcript levels for *gloA* and *gloB* using RT-qPCR, and found that COH1 has higher abundance of both *gloA* and *gloB* transcripts compared to CJB111 and A909 (**Fig. S2F**). Altogether, the predicted GloA protein in GBS contains conserved residues important for enzyme activity and metal binding but it is not clear if the common A45S amino acid change correlates with enzyme activity. In addition, transcript expression for *gloA* and *gloB* are increased in COH1 which could explain higher MG resistance.

**Figure 2.**
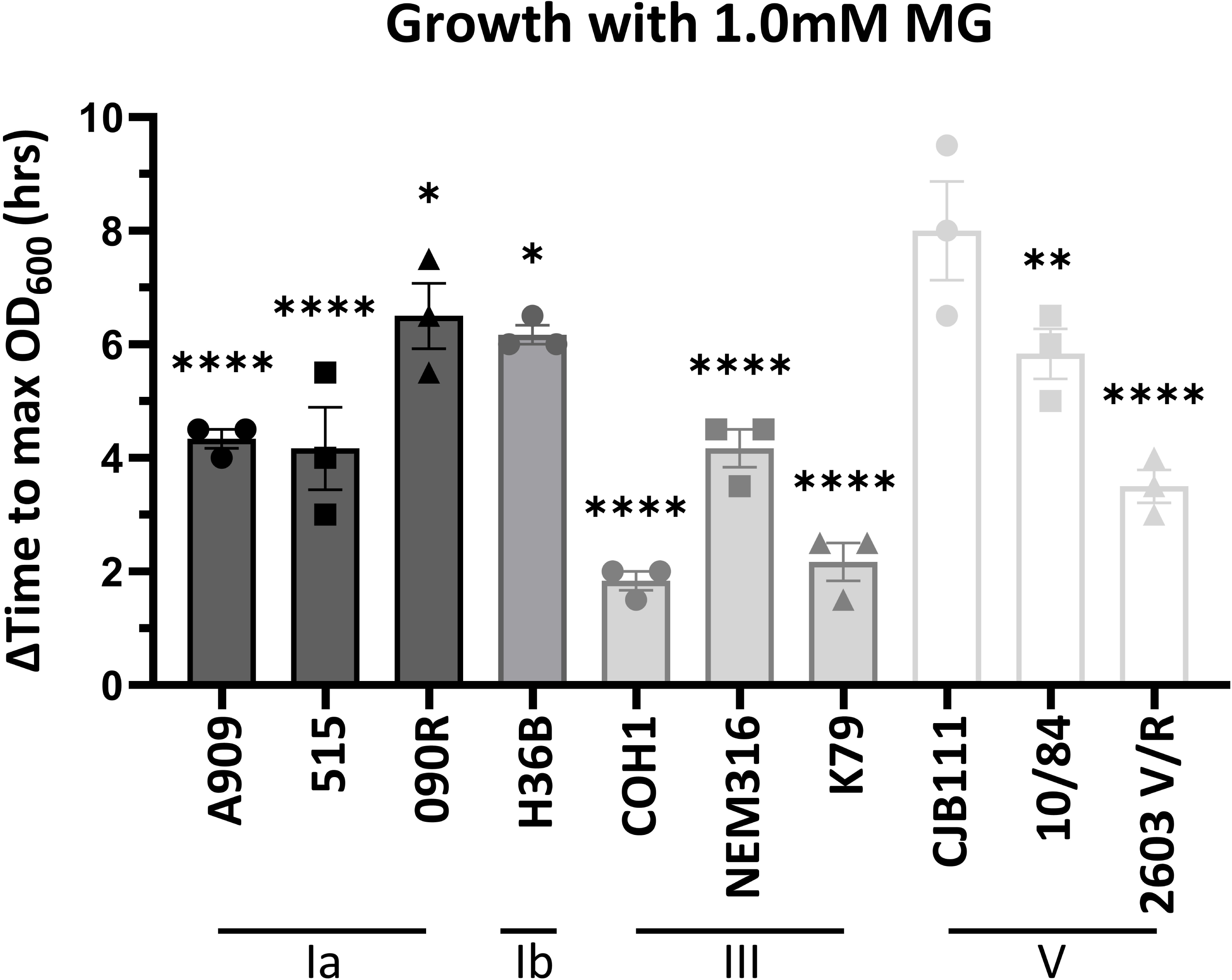
MG sensitivity differs across GBS isolates. Change in time to max OD_600_ for growth curves with 1.0 mM MG compared to no MG in mCDM for representative serotype Ia, Ib, III, and V GBS strains. Significance determined by comparing all strains to the most sensitive strain, CJB111, by one-way ANOVA with uncorrected Fisher’s LSD test with *P* < 0.05. * < 0.05, **<0.01, **** < 0.0001.

### Glyoxalase A contributes to methylglyoxal detoxification in GBS

To confirm our Tn-sequencing results we chose to study the first enzyme in the glyoxalase pathway, GloA. Using allelic exchange mutagenesis, we constructed a mutant in *gloA* (Δ*gloA*) and a complemented strain (p*gloA*) in CJB111 as described in Materials and Methods. MG detoxification was then tested using these strains by MG quantification and growth curve analysis. First, to measure if MG accumulates in the Δ*gloA* strain, we measured MG concentrations using an ELISA on lysed cell pellet samples for CJB111 WT, Δ*gloA*, and p*gloA* strains. The concentration of MG was normalized to the total protein concentration of each sample and found to be significantly increased in the Δ*gloA* mutant compared to the CJB111 WT and complemented strains (**Fig. 3A**). To determine if this accumulated MG in the Δ*gloA* mutant may be toxic/impact GBS growth, all strains were inoculated into mCDM with or without the addition of 0.5 mM MG. Indeed, a greater growth delay was observed for Δ*gloA* with a change in time to max OD_600_ at 4.92 hrs compared to 3.58 hrs for WT or 2.83 hrs for p*gloA*, which confirms *gloA* is involved in MG detoxification (**Fig. 3B**). Furthermore, the OD_600_ at 8 hrs was compared between strains and confirmed a significant decrease in Δ*gloA* growth compared to WT or the complemented strain following the addition of MG (**Fig. 3C**). To determine if the growth delay we observed with MG was due to the an emerging resistant subpopulation, we grew CJB111 WT to mid-logarithmic phase with 0.5 mM MG and then inoculated fresh media with or without increasing MG (**Fig. S3A**). From this experiment we still observed a growth delay with fresh MG indicating it is not a more resistant subpopulation emerging. As MG is primarily produced from glycolysis in cells (bacteria and host), we further investigated the impact of GloA on GBS growth in mCDM with increasing glucose concentrations. However, we did not observe a growth defect for the Δ*gloA* mutant compared to WT or complemented strain at low (biologically relevant blood glucose concentration), medium (ideal concentration for GBS growth), or high glucose concentrations tested (**Fig. S3B**). Additionally, upon assessment of general virulence characteristics, we also did not observe a difference in susceptibility to hydrogen peroxide or hemolytic activity between CJB111, Δ*gloA*, and p*gloA* (**Fig. S3C&D**). Taken together MG quantification and growth analysis suggest that GloA contributes to MG detoxification in GBS. Our results also suggest that, under the conditions tested, GBS may not produce enough MG from glucose metabolism to negatively impact its own growth.

**Figure 3.**
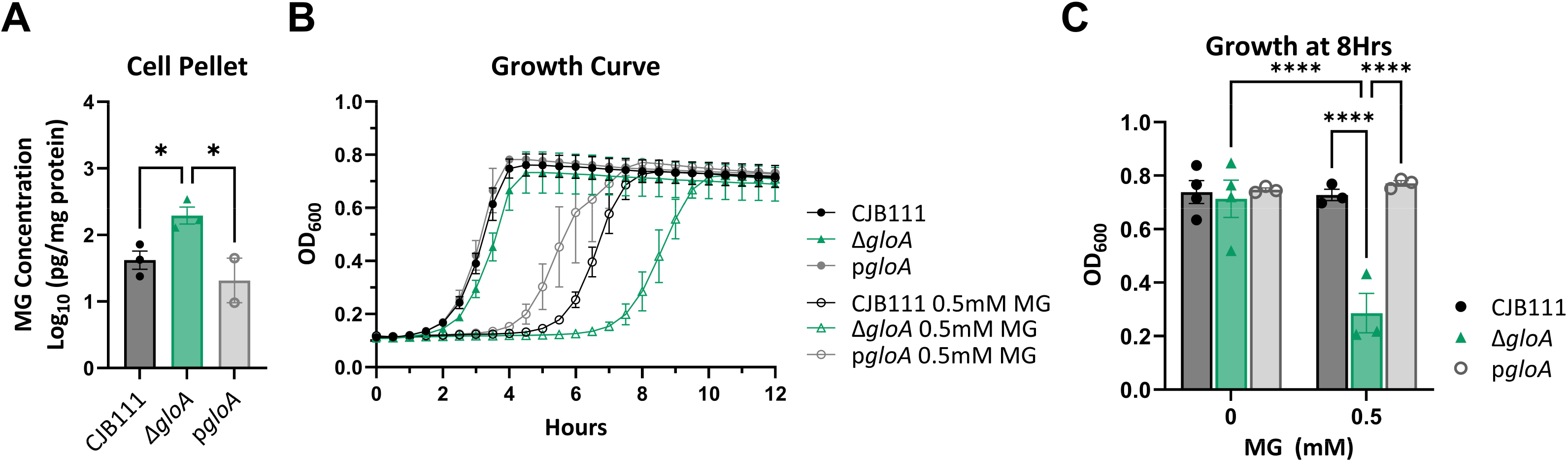
The contribution of glyoxalase A to GBS methylglyoxal detoxification. (A) ELISA MG quantification of cell pellets for WT CJB111, Δ*gloA*, and p*gloA* strains after growth in mCDM for 4 hours. (B) Growth curve measured by OD_600_ for WT CJB111, Δ*gloA*, and p*gloA* strains grown with or without 0.5 mM MG in mCDM. (C) Comparison of growth shown in (B) at 8hrs between strains. Significance determined by (A) One-way ANOVA or (C) 2way ANOVA with Fisher’s LSD multiple comparisons test with *P* < 0.05. * < 0.05, **** < 0.0001.

### Glyoxalase A is necessary for GBS survival *in vivo*

To further confirm the Tn-sequencing results and determine if *gloA* is important during infection, we repeated our bacteremia model of infection by intravenously injecting mice with 1.5-2 x 10^7^ CFU CJB111 WT or the Δ*gloA* mutant and monitoring the infection for up to 72 hours post-infection. Mice infected with the Δ*gloA* mutant exhibited significantly decreased mortality compared to those infected with WT, with greater than 75% surviving to the experiment endpoint (**Fig. 4A**). In order to monitor CFU burden over-time, blood samples were taken from surviving mice at 24 and 48 hrs post-infection and at the TOD (**Fig. 4B**). Mice infected with Δ*gloA* had significantly decreased blood burdens as soon as 24 hrs post-infection and in tissue burdens at the time of death, indicating the mutant strain is not able to survive as well in the bloodstream and disseminate to other organs compared to WT CJB111 (**Fig. 4B&C**). It is important to note that all of the strains had similar blood burdens at 6 hrs post-infection (**Fig. S4A**) and that the Δ*gloA* infected mice that succumbed to the infection before 72 hours had the highest CFU counts in the blood at 24 hours (**Fig. 4B**). Additionally, we infected mice with the complemented strain and confirmed that it was able to survive in the blood longer than Δ*gloA* (**Fig. S4**).

**Figure 4.**
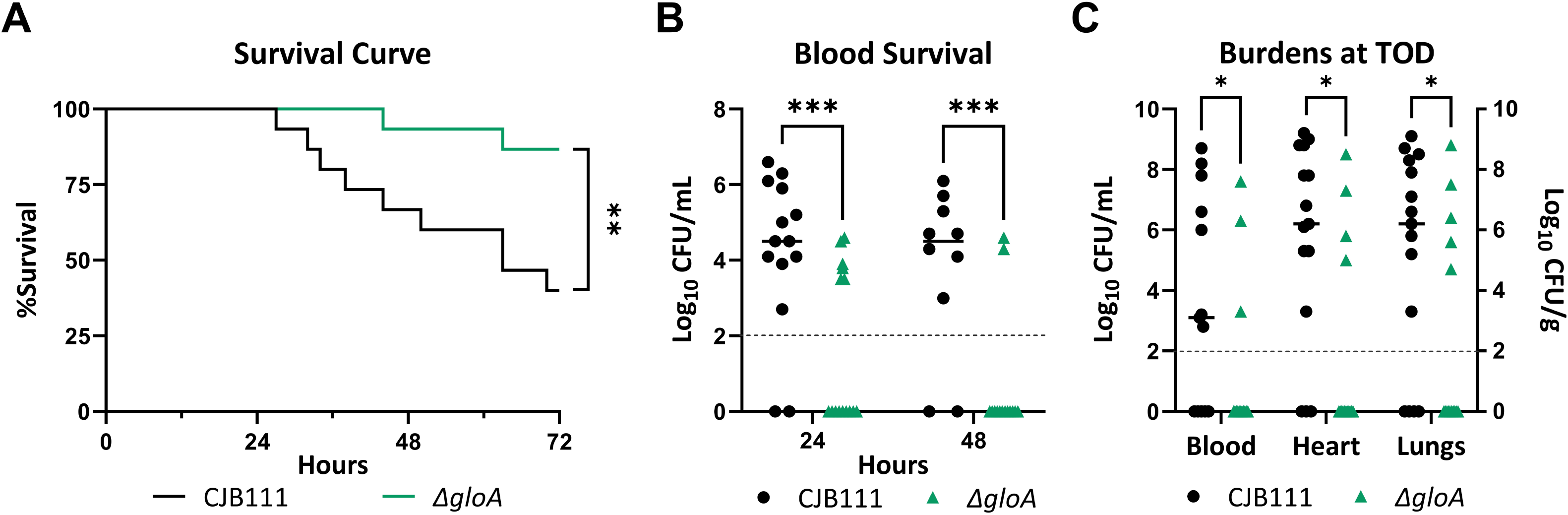
Methylglyoxal detoxification is necessary for GBS infection. (A) Survival curve of mice tail-vein injected with 10^7^ CFU WT CJB111 or Δ*gloA.* (B) Recovered CFU counts from the blood of infected mice at 24 and 48 hours post-infection. (C) Recovered CFU counts from the blood, heart, and lungs of infected mice at time-of-death. In B&C, limit of detection is indicated with dotted line. Significance determined by (A) Log-rank (Mantel-Cox) test, (B) Mixed-effects analysis with uncorrected Fisher’s LSD test, or (C) multiple Mann-Whitney *U* tests with *P* < 0.05. * < 0.05, ** < 0.01, ***<0.001.

### Importance of GBS Glyoxalase A to Neutrophil Survival

Pro-inflammatory M1-type macrophages primarily use glycolysis to generate energy (*68*) and have been shown to produce aldehydes, such as MG, in response to infection (*66, 69–71*). However, neutrophils are the primary immune cell GBS encounters during acute infection (*55*), which also utilize glycolysis as their primary energy source (*66, 68–72*). Therefore, to determine if GBS *gloA* contributes to neutrophil survival we performed *in vitro* neutrophil killing assays using differentiated HL60-neutrophils with the CJB111 WT, Δ*gloA*, and p*gloA* strains. At 5 hrs post-infection, the Δ*gloA* mutant strain exhibited significantly decreased survival compared to WT or the complemented strains (**Fig. 5A**). This phenotype was independent of serum killing and cytotoxicity since serum by itself did not impact GBS survival over time and there were low levels and no differences in HL60 cytotoxicity between the strains at 5 hrs (**Fig. S5A&B**). We also confirmed that Δ*gloA* has decreased survival in the presence of primary murine bone marrow neutrophils (BMNs) (**Fig. S5C**). Next, we used cytochalasin D treatment to block phagocytosis of HL60-neutrophils which resulted in a significant increase in survival for CJB111 WT and p*gloA* strains (**Fig. 5B**). The Δ*gloA* mutant also had a minor increase in survival with cytochalasin D treatment but it was not significant when compared to the DMSO control and there was still a significant decrease in survival when compared to WT and complement indicating the mutant is susceptible to multiple mechanisms of neutrophil killing. To evaluate if this increased killing might be due to a general increase in neutrophil production of MG in response to GBS infection, we measured the accumulation of intracellular MG-modified proteins in HL60-neutrophils using flow cytometry. Consistent with the literature (*30*) we observed that all cells contained detectable MG-modified proteins. Interestingly, however, upon infection, we observed a significant increase in anti-MG geometric mean fluorescent intensity (MFI) compared to uninfected controls (**Fig. 5C**), indicating that infection increases intracellular MG within HL60s. This increase was also only observed in high glucose conditions (**Fig. S5D**), which is similar to what has been described for *S. pyogenes* where they reported that GloA contributed to survival against neutrophils with elevated glucose (*65*). To examine the contribution of neutrophils controlling GBS infection *in vivo,* we depleted neutrophils in mice prior to intravenous infection. Mice were injected intraperitoneally with anti-Ly6G or an isotype control 24 hrs (*73*) before intravenous infection with 1 x 10^7^ CFU CJB111 or Δ*gloA*. Upon assessing morbidity and mortality of these groups over 72 hours post-infection, we observed that both CJB111 and Δ*gloA*-infected neutrophil depleted mice exhibited significant increase in mortality when compared to their non-depleted cohorts (**Fig. 5D**), although percent survival of neutrophil-depleted mice infected with Δ*gloA* remained higher than neutrophil-depleted mice infected with CJB111. At 12 hours post-infection, we observed that neutrophil depletion abrogated the attenuated phenotype of the Δ*gloA* mutant in the blood, as the CJB111 and Δ*gloA*-infected, neutrophil depleted mice did not differ in blood burdens (**Fig. 5E**). Further, CJB111 and Δ*gloA* CFU burdens were significantly increased in the neutrophil-depleted mice compared to the non-depleted mice. Taken together these results show that the attenuation of the Δ*gloA* mutant can be partially rescued with neutrophil depletion and suggest that MG produced by other cell types may aid in the defense against GBS bloodstream infections.

**Figure 5.**
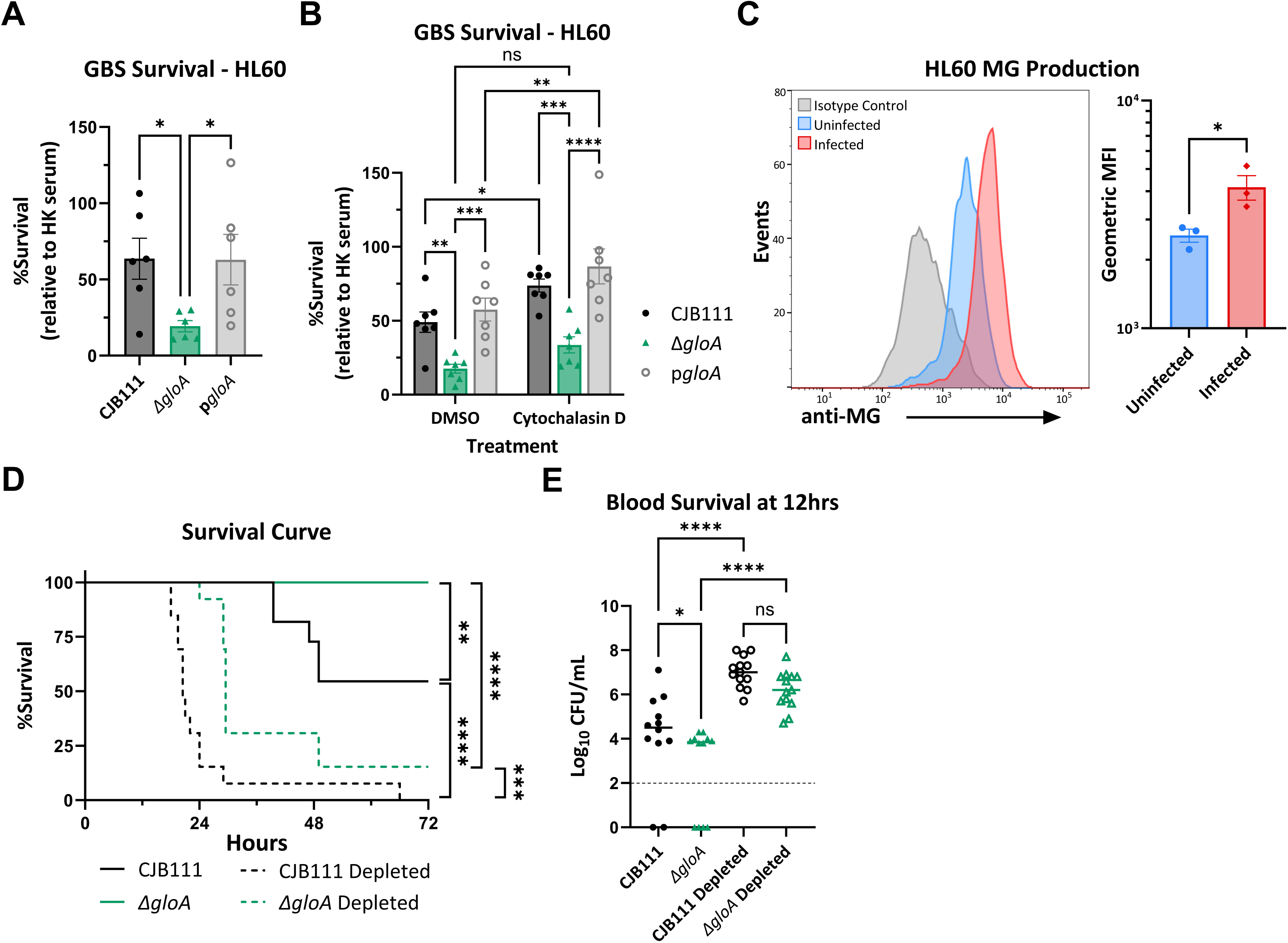
Methylglyoxal detoxification is necessary for GBS survival against neutrophils. Survival of WT CJB111, Δ*gloA*, and p*gloA* strains after 5 hours infection of HL60-neutrophils (A) without treatment or (B) with DMSO control or Cytochalasin D treatment. (C) Flow cytometry quantification of intracellular MG-modified proteins in HL60-neutrophils with or without WT CJB111 infection. Left: Representative histogram displaying MG signal that includes isotype control, uninfected, and infected cells. Right: Geometric mean fluorescent intensity (MFI) quantification for MG in uninfected and infected samples. (D) Survival curve of normal or neutrophil depleted mice that were tail-vein injected with 10^7^ CFU WT CJB111 Δ*gloA*. (E) Recovered CFU counts from the blood of infected mice 12 hours post-infection. Limit of detection is indicated with dotted line. Significance determined by (A&E) One-way ANOVA with Fisher’s LSD multiple comparisons test. (B) Two-way ANOVA with Fishers LSD multiple comparisons test. (C) Unpaired Student *t* test, (D) Log-rank (Mantel-Cox) test with *P* < 0.05. * < 0.05, **<0.01, *** < 0.001, **** < 0.0001.

## Discussion

GBS must be able to survive multiple host niches to cause invasive infection in neonates. Some of these environments include the vaginal tract, amniotic fluid, and blood (*1, 74*). Tn-sequencing is a powerful and common method for investigating bacterial genes necessary for survival and fitness in these different environments. Previously, an *ex vivo* Tn-sequencing was performed in human blood by Hooven *et al.* 2017 using a TnSeq library in the GBS A909 serotype Ia background (*75*). Their results found similar underrepresented genes compared to our *in vivo* dataset such as genes involved in capsule biosynthesis, metal homeostasis, and arginine metabolism. Interestingly, they identified *relA* to be underrepresented, which encodes a GTP pyrophosphokinase and is a central regulator of the stringent response in GBS. They found that *relA* not only controls stringent response activation and the arginine deiminase pathway but also impacts βH/C production. While we did not observe *relA* in our dataset, other putative stress response proteins like *ytgB* and *asp1* were significantly underrepresented. Most notably, *asp1* is annotated as an Asp^23^/Gls^24^ family envelope stress response protein and was found to be upregulated when GBS was incubated in human blood (*13, 76*) and downregulated after exposure to high glucose (*77*). We also observed that the two-component system (TCS) *dltRS* and a *dlt* gene were underrepresented, which are involved in modulating surface charge and contribute to cationic antimicrobial peptide resistance and decrease phagocytic killing (*35, 54*).

Previous studies have shown that a *dltA* mutant exhibited decreased virulence in a murine model with significantly lower burdens in tissue and blood compared to WT GBS (*35*). Another TCS, *bceRS/nsrRK,* exhibited the highest negative fold change of the underrepresented TCS and has been shown to contribute to bacitracin (antibiotic), nisin (lantibiotic), cathelicidin/LL-37 (human antimicrobial peptide), and oxidative stress resistance (*56, 57*). Its role in GBS pathogenesis was demonstrated as a *bceR* mutant yielded decreased virulence during murine infection and decreased biofilm formation (*56*). Another top gene hit identified within the present study to be important for GBS blood survival is the C5a peptidase *scpB. scpB* is already known to be involved in complement evasion and fibronectin binding and is associated with neonatal isolates (*36, 78*). Overall, identifying these known virulence factors in our study demonstrates the validity of our *in vivo* Tn-sequencing screen to identify novel factors important for blood survival in mice and supports previous research in the streptococcal field. It also provides another resource for developing new hypotheses and research projects. For example, methylglyoxal (MG) detoxification has not been previously characterized in prior GBS studies.

MG is a highly reactive electrophilic species (RES) and byproduct of normal cell metabolism which can be spontaneously or enzymatically produced by all cells (*28, 30*) with up to 90% of cell MG estimated to come from glycolysis alone (*79*). Notably, MG is also a precursor to advanced glycation end products (AGEs) and is associated with many other human diseases like diabetes, cancer, and neurological disorders like Alzheimer’s disease (*30, 80*). The most well-known and ubiquitous pathway for MG detoxification is the glyoxalase pathway which consists of glyoxalase A (*gloA*) and glyoxalase B (*gloB*) enzymes. Recently, the glyoxalase pathway, especially *gloA*, in *L. monocytogenes* was found to contribute to intracellular survival in macrophages and during murine infection (*66*). In addition, the glyoxalase pathway in *S. pyogenes* was shown to be important for survival against neutrophils in a glucose and myeloperoxidase dependent manner (*65*). The GBS *gloA* and *gloB* homologs were underrepresented in our Tn-sequencing dataset therefore, we hypothesized that GBS may encounter host-derived MG during bacteremia as a response to infection (*65, 69, 81, 82*). The first step in the glyoxalase pathway is mediated by GloA and, in this study, we have characterized its contribution to GBS MG tolerance *in vitro* and infection *in vivo*. We observed that a Δ*gloA* mutant exhibited decreased survival in the presence of HL60-neutrophils as well as primary murine neutrophils and that its virulence was largely restored in neutrophil-depleted mice. We also measured an increase in MG-modified proteins in HL60-neutrophils upon GBS infection which is likely from increased production of MG by the cells themselves. These results indicate MG-mediated killing may constitute another important defense mechanism used by immune cells to kill invading bacteria.

We also observed that tolerance to MG varies across GBS isolates and is not definitively correlated to serotype, GloA amino acid sequence or glyoxalase gene regulation. Of note, two of the serotype III strains, the most commonly isolated serotype from neonatal invasive infections (*83*), had the first and second highest overall tolerance to MG, but additional strains would need to be tested to determine if serotype III have higher average MG tolerance. Interestingly, COH1 (serotype III), which had the highest resistance to MG, was the only strain tested that had the A45S variant in GloA and higher baseline transcript levels of *gloA* and *gloB* compared to CJB111 and A909 strains. The A45S variant is also only 10 amino acids away from the K55 residue which is predicted to be involved in metal binding and could impact folding or metal coordination (*67, 84, 85*). However, other GBS strains tested also had high resistance to MG, like K79, which does not have the A45S variant. Therefore, it is unlikely that this amino acid change is the sole determinant of enzyme activity, but further investigation is needed to determine if GloA protein variants and regulation impact GBS MG tolerance. Relatedly, *S. mutans* was shown to be more tolerant to MG than most other commensal oral Streptococcal species and was also shown to outcompete *S. sanguinis* when MG was present in a competition experiment (*86*). From this previous study and our work shown here, we hypothesize that differences in GBS MG tolerance could be influenced by the environments they were found in, like the presence of other bacterial species or host differences.

Components of metal transport systems were also significantly underrepresented in the Tn-sequencing dataset with top hits including the zinc import system *adcAAIIBC* and *lmb,* the manganese import system *mtsABC,* and the putative nickel import system *nikABCDE* (**Table 1**). These results suggest GBS requires trace metals to survive in the blood. Previously it has already been shown previously that zinc and manganese transporters are important for maintaining GBS metal homeostasis and contribute to vaginal colonization and female reproductive tract ascension as well as blood survival (*13, 32, 76, 87*). In addition, both zinc and manganese import systems are important in combating nutritional immunity, or the sequestration of nutrients by the host, mediated by a neutrophil-produced metal-binding protein called calprotectin (*33, 46*). The nickel transporter, however, has not been as well characterized. In our previous study, we attempted to measure nickel concentrations in a *nikA* mutant, but it was under the limit of detection in our samples. However, we did observe lower levels of copper indicating the system could be transporting more than one metal (*47*). There are only 9 known enzymes present in archaea, bacteria, plants, and primitive eukaryotes that are nickel dependent, with urease being the most notorious, however, GBS does not encode a urease enzyme (*32, 47, 88, 89*). Interestingly, another of the 9 known nickel-dependent enzymes is GloA which was found to use nickel (Ni^2+^) as a cofactor in *E. coli* (*34, 67, 85*). Therefore, the importance of the nickel transporter in the blood could be due to increased GloA activity but the requirement for nickel in GBS still remains to be elucidated.

MG is formed primarily from glycolysis but it can be produced, albeit to a lesser extent, from lipid, ketone, and protein metabolism (*30, 34*). MG is toxic to cells due to its electrophilic properties allowing it to react with different molecules, like DNA and protein, and effectively arrest growth (*81*). For example, MG has been shown to increase mutation rates in *L. monocytogenes* by binding DNA (*66*) and inhibits protein synthesis and modification by binding to guanine and arginine residues (*90–93*). It is important to note that in some bacteria, like *E. coli* and *L. monocytogenes,* MG can be formed directly from dihydroxyacetone phosphate (DHAP) during glycolysis by methylglyoxal synthase (*mgsA*); however, GBS, like other streptococci, do not have a MG synthase gene (*65, 86*). The lack of a synthase gene further supports our hypothesis that GBS encounters host-derived MG toxicity during infection. Since approximately 99% of cellular MG is thought to be already bound to molecules like DNA and protein it is difficult to quantify accurately; however, intracellular MG concentrations are consistently estimated below 10 µM and are known to be dependent on glutathione concentrations (*30, 94, 95*). In serum of diabetic individuals, the concentration of MG and MG-derived AGEs are increased compared to healthy individuals, most likely due to increased glucose concentrations (*95*). MG is historically tied to diabetes because it is known to exacerbate diabetic complications like microvascular and kidney dysfunction and contribute to the progression of the disease (*95*). Our lab has shown that GBS is a common colonizer of infected diabetic wounds (*96*) and we show here the production of MG-modified proteins by HL60-neutrophils is dependent on glucose and GBS infection (**Fig. 5C**, **Fig. S5D**). Therefore, research into the role of the GBS glyoxalase pathway in the context of diabetic wound infection is a current area of study.

Lastly, conversion of MG to D-lactate by the glyoxalase pathway was first described over 100 years ago and is the most ubiquitous and conserved process for MG detoxification across all kingdoms of life (*28*). MG detoxification has not been thoroughly studied in streptococci in the context of disease and has never been characterized in GBS. Thus far, studies focusing on *S. pyogenes*, *S. mutans*, and *S. sanguinis* have shown *gloA* to be the primary modulator of MG tolerance *in vitro* with *gloB* mutants having little effect (*65, 86*). This is in support of what was observed with *L. monocytogenes,* but not *Salmonella* where it was found that *gloB* was more important for *Salmonella* resistance to oxidative stress and killing by macrophages (*66, 97*). Here, we found *gloA* to be dispensable to GBS tolerance of H_2_O_2_ (**Fig. S3C**) but the contribution of *gloB* remains to be examined. Additional enzymes known to break down MG into acetol or lactaldehyde intermediates include aldose, aldehyde, and MG reductases. A putative aldo/keto reductase (*yvgN*) was significantly underrepresented in our data set (**Table 1**) but its contribution to GBS virulence requires further investigation (*34*).

In this study we demonstrate, for the first time, an essential role of the glyoxalase pathway to GBS MG resistance and overall pathogenicity during blood stream infection. We investigated the role of GloA *in vitro* and *in vivo* and confirmed it is important for growth in the presence of MG, survival against neutrophils, and during invasive infection. Our study also provides further evidence in support of the aldehyde hypothesis in that MG detoxification is an important component for bacterial survival against host defenses; however, the role of the glyoxalase system to GBS survival in macrophages required further investigation (69). Specifically, we show increased MG production by neutrophils in response to infection. Overall, research aimed at understanding metabolic mechanisms used by bacteria to survive in the blood and RES toxicity will be important for the development of new treatment and therapies for infection and will expand our knowledge about host-pathogen interactions.

## Materials and Methods

### Bacterial strains, media, and growth conditions

See Table S2 for strains and primers used in this study. GBS strains were grown statically at 37°C in THB, unless otherwise stated. Streptococcal chemically defined medium (*64*) was modified by omitting L-cysteine and adding 22mM glucose, unless otherwise stated. *Escherichia coli* strains for cloning were grown in LB at 30°C or 37°C with rotation at 250 rpm. Kanamycin and erythromycin (Sigma-Aldrich, St. Louis, MO) were supplemented to media at 50 µg/mL and 500 µg/mL, respectively, for *E. coli*. Kanamycin, spectinomycin, and erythromycin (Sigma-Aldrich, St. Louis, MO) were supplemented to media at 500 µg/mL, 100 µg/mL, and 5 µg/mL, respectively, for streptococcal strains.

### Routine molecular biology techniques

All PCR reactions utilized Phusion or Q5 polymerase (Thermo Fisher, Waltham, MA). PCR products and restriction digest products were purified using QIAquick PCR purification kit (Qiagen, Venlo, NL) per manufacturer protocols. Plasmids were extracted using QIAprep miniprep kit or plasmid midi kit (Qiagen, Venlo, NL) per manufacturer protocols. Restriction enzyme digests utilized XmaI, EcoR1, and BamH1 (New England Biolabs, Ipswich, MA) for 2 hours at 37°C in a thermocycler. Ligations utilized Quick ligase (New England Biolabs, Ipswich, MA) at room temperature for 5 min or Gibson Assembly Master Mix (New England Biolabs, Ipswich, MA) per manufacturer protocols. All plasmid constructs were sequence confirmed by Sanger sequencing (CU Anschutz Molecular Biology Core, Aurora, CO) or whole plasmid sequencing (Quantara Biosciences, Hayward, CA).

The mutant strains were generated as previously described (*12, 32*). Briefly, for the *gloA* mutant, genomic 5’ and 3’ regions flanking the *gloA* gene were amplified and fused with a spectinomycin cassette by FailSafe PCR (Lucigen, Middleton, WI). Fragments and pHY304 vector were digested with restriction enzymes and ligated using Quick Ligase. The ligation reaction product was transformed into chemically competent *E. coli*. pHY304 plasmids were purified from *E. coli* and electroporated into GBS CJB111 genetic background. Constructs were confirmed by PCR and sequencing. Complement strains were generated by amplifying the *gloA* gene in GBS and linearizing pABG5 by PCR. Products were ligated using Gibson assembly and then transformed into chemically competent *E. coli*. Plasmids were purified from *E. coli* and electroporated into GBS CJB111 Δ*gloA* genetic background. Primers used in the construction of strains are listed in Table S2. The mutant had no growth or hemolysis defects observed (**Fig. S3B&D**).

### Study approval

Animal experiments were approved by the Institutional Animal Care and Use Committee (IACUC) at the University of Colorado Anschutz Medical Campus protocol #00316 and were performed using accepted veterinary standards. The University of Colorado Anschutz Medical Campus is AAALAC accredited; and its facilities meet and adhere to the standards in the “Guide for the Care and Use of Laboratory Animals”. All mice were purchased from Charles River Laboratories (CD1) and housed in pathogen-free, biosafety level-2 animal facilities.

### *In vivo* transposon screening

Triplicate cultures of the pooled CJB111 pKrmit transposon library (*32*) were grown overnight at 37°C in THB with kanamycin at 300 µg/mL and back diluted to an OD600 0.4. Libraries were normalized to ∼4 x 10^7^ CFU/100 µL and injected via tail-vein into 6–8-week-old CD-1 male mice using the established hematogenous infection model (*98–101*). Blood was collected by cardiac puncture between ∼18-28 hours post infection. 100 µL of input library and blood was plated in duplicated on CHROMagar Strep B with 300 µg/mL kanamycin and incubated overnight at 37°C to collect recovered transposon mutants. Bacterial growth from spread plates were collected and 3-4 mice per library were pooled together, genomic DNA extracted using ZymoBiomics DNA miniprep Kit (Zymo Research).

### Transposon library sequencing

Libraries were prepared and sequenced at the University of Minnesota Genomics Center (UMGC) according to https://www.protocols.io/view/transposon-insertion-sequencing-tn-seq-library-pre-rm7vzn6d5vx1/v1. Briefly, genomic DNA was enzymatically fragmented, and adapters added using the NEB Ultra II FS kit (New England Biolabs), and ∼50 ng of fragmented adapted gDNA was used as a template for enrichment by PCR (16 cycles) for the transposon insertions using mariner-specific (TCGTCGGCAGCGTCAGATGTGTATAAGAGACAGCCGGGGACTTATCATCCAACC) and Illumina P7 primers. The enriched PCR products were diluted to 1ng/ul and 10 ul was used as a template for an indexing PCR (9 cycles) using Nextera_R1 (iP5) and Nextera_R2 (iP7) primers. Sequencing was performed using 150-base paired-end format on an Illumina NextSeq 2000 and Illumina NovaSeq 6000 system to generate ∼40-60 million reads per library.

### Tn-sequencing analysis

The R1 reads from both sequencing runs were concatenated and quality was assessed using FastQC (*102*)(http://www.bioinformatics.babraham.ac.uk/projects/fastqc/). Reads were trimmed using Cutadapt (v 4.2) (*103*) with the following parameters; sequence length with a minimum of 12 bases, removal of flanking “N” bases, reads were trimmed of 3’ “G” bases, and reads were trimmed with the reverse complemented mariner transposon sequence (ACTTATCAGCCAACCTGTTA). TRANSIT (v 3.2.7) (*104*) was used to align trimmed reads to the CJB111 genome (CP063198) and for analysis of transposon insertion sites. The Transit PreProcessor processed reads using default parameters with the Sassetti protocol, no primer sequence, and mapped to the genome sequence using Burrows-Wheeler Alignment (BWA) (*105*). Insertion sites were normalized using the Total Trimmed Reads (TTR) method in TRANSIT and analyzed using the resampling method to compare the insertion counts recovered in blood vs the input library using default parameters, with the addition of ignoring TA sites within 5% of the 5’ and 3’ end of the gene. Significance determined by p_adj_ <0.05 and log_2_FC <-1 or >1. All sequencing reads have been deposited into NCBI SRA under BioProject ID PRJNA1125445.

### Murine model of bloodstream infection

We infected mice as previously described for CJB111 to cause invasive infection (*98–101*). Briefly, 15 8-week-old CD1 male mice were intravenously challenged with 1.5-2 x 10^7^ CFU GBS CJB111, Δ*gloA,* or p*gloA*. At 6, 12, 24, and/or 48 h post-infection, blood samples were taken by tail prick and plated on THA to quantify GBS CFU burden. Once mice reached moribund state or 72 h post-infection mice were sacrificed, and blood was harvested by cardiac puncture and lung and heart tissue were removed and homogenized in sterile PBS. All samples were plated on THA or CHROMagar to quantify GBS CFU burden. For neutrophil depletion, 11-13 mice per group were given 200µg *InVivo*MAb anti-mouse Ly6G antibody (Bio X Cell, Lebanon, NH) or 200µg *InVivo*MAb rat IgG2a isotype control diluted in *InVivo*Pure pH 7.0 dilution buffer by intraperitoneal injection 24 h before infection.

### GloA protein comparisons

GloA amino acid sequences where aligned using ClustalOmega (*106*) and the alignment figure was created using the ESPript Server (*107*)(https://espript.ibcp.fr). Protein IDs used: ABA45143.1 (GBS A909), QOW77196.1 (GBS CJB111), WP_001116201.1 (GBS COH1), WP_002985686.1 (GAS 5448) and P0AC81.1 (*E. coli* K-12). GBS GloA phylogenetic tree was generated using NCBI BlastP (*108, 109*) and visualized using FigTree v1.4.4 (http://tree.bio.ed.ac.uk/software/figtree/). The protein ID QOW77196.1 (GBS CJB111) was used as the query against *S. agalactiae* and only proteins with percent identity and query cover greater than 50% are shown. The dimeric structure of the *S. agalactiae* GloA was predicted using AlphaFold2 (*110*) as implemented in ColabFold (*111*). PyMOL (version 2.5.2, Schrödinger, LLC.) was used to create images of the predicted GloA structure and the *E. coli* glyoxalase I crystal structure RCSB PDB entry 19FZ (*67, 112*) (RCSB.org).

### *In vitro* growth comparisons

Overnight cultures of Streptococcal strains were diluted 1:100 in mCDM with or without methylglyoxal (MG, Sigma-Aldrich M0252, St. Louis, MO) or hydrogen peroxide 3% w/w (VWR, Radnor, PA) at concentrations listed in figure legends in a 96-well plate. For strain/serotype growth comparisons with MG, the overnight cultures were first normalized to OD600 0.6 in PBS before being diluted to 1:100 in mCDM with or without MG. For testing the effects of MG pre-exposure on growth, overnight cultures were diluted 1:100 in mCDM with 0.5mM MG and allowed to grow until mid-logarithmic phase (OD600 0.4-0.6). Then the mid-logarithmic cultures were used to inoculate a new 96-well plate with fresh mCDM with or without MG. For all growth curves longer than 8 hrs the plate was covered with a Breathe-Easy gas permeable sealing membrane (USA Scientific, Ocala, FL) and then incubated at 37°C without shaking in a Tecan Infinite M Plex for up to 24 hrs with OD600 taken every 30 min. For growth curves with CFU/mL shown, the 96-well plate was incubated at 37°C without shaking and samples were taken every 2 hrs for dilution plating on THA. All growth curves were performed in biological triplicate.

### ELISAs on culture pellets

Overnight cultures of Streptococcal strains were diluted 1:100 in mCDM and then grown for 4 h at 37°C. 3 mL of each culture was pelleted, re-suspended in PBS, and then homogenized using 0.1 mm dia. Zirconia beads. Methylglyoxal concentration in culture samples was measured using an ELISA kit (Biomatik EKN53482, Kitchener, ON, CA) per manufacture instructions. BSA protein assay standard was used to quantify protein concentration in each culture sample and each group was performed in biological duplicate or triplicate.

### RT-qPCR

Samples were made by centrifuging 1 mL aliquots of cultures grown to mid-log phase in mCDM and then re-suspending in 1 mL fresh mCDM and incubating for 30 min at 37°C. 1 mL of RNAProtect Bacteria Reagent (Qiagen, Venlo, NL) was then added before centrifuging and washing pellets with ice cold PBS. Sample RNA was prepped using the NucleoSpin RNA kit (Macherey-Nagel, Dueren, DE) and TURBO DNase treated (Invitrogen by Thermo Fisher, Waltham, MA) per manufacture instructions. 250 ng of RNA was made into cDNA for each sample using the qScript cDNA Synthesis Kit (Quantabio, Beverly, MA) per manufacture instructions. cDNA was then diluted 1:20 in water and RT-qPCR run using PerfeCTa SYBR Green FastMix (Quantabio, Beverly, MA) per manufacture instructions and *glcK*, *gloA*, and *gloB* qPCR primers (see Table S2). Each sample was run in technical duplicate for each gene. Each sample Cq value for *glcK*, *gloA*, and *gloB* was normalized to the total average CJB111 Cq value for each gene, respectively.

### Hemolysis assay

Overnight cultures of Streptococcal strains were diluted 1:100 in THB and then grown to mid-log phase at 37°C. Cultures were then normalized to OD600 0.4 in PBS. 400 µL blood, 400 µL PBS, and 200 µL normalized culture was added to each sample microfuge tube. 400 µL blood and 600 µL sterile water was added to positive control tubes and 400 µL blood and 600 µL PBS was added to negative control tubes. Tubes were made in technical duplicate and incubated at 37°C with rotation. At 24 hrs, 100 µL aliquots were taken and centrifuged at 5500 x g for 1 min. OD543 of supernatant was measured using a Tecan Infinite M Plex and %lysis calculated by subtracting negative control from all samples and then dividing samples by positive controls.

### HL60-neutrophil and primary bone marrow neutrophil killing assays

HL60 cells were cultured in RPMI + 10% FBS, differentiated with 1.25% DMSO (Sigma-Aldrich, St. Louis, MO) for 4 days, and infected as previously described (*113*). Briefly, GBS strains were grown to mid-log, normalized in PBS, and then opsonized in 10% normal human serum or heat killed (HK) serum in FBB buffer (0.5% BSA and 2.2mM CaCl_2_ in HBSS) for 15 min in a 96-well plate. HL60 cells were diluted to desired concentration in FBB buffer and then either pre-incubated with 20 µM cytochalasin D (Sigma-Aldrich C8273, St. Louis, MO) or DMSO (Sigma-Aldrich, St. Louis, MO) vehicle control for 15 min with rotation or immediately added to each well with opsonized bacteria to reach an MOI of 0.002. The plate was incubated at 37°C with shaking for 5 hrs. %Survival at 5 hrs was calculated by dividing CFU recovered from wells with GBS opsonized with normal serum by CFU recovered from wells with GBS opsonized with heat-killed (HK) serum. CFU from control wells without HL60 cells were also quantified.

Murine neutrophils were isolated from bone marrow of 8-12-week-old C57B6/J mice using an Anti-Ly-6G Microbeads kit (Miltenyi Biotec 130-120-337, Germany) per the manufacturer’s instructions. Neutrophil isolations yielded ≥ 95% purity and were used within 1 h of isolation. Murine bone marrow neutrophils (BMNs) were re-suspended in RPMI + 5% FBS then adhered to TC-treated 96-well plates for 30 min prior to infection. Neutrophils were infected at an MOI of 0.01 and then synchronized by centrifugation (200 x g for 5 min). Neutrophils were incubated for 4 h in a cell culture incubator at 37°C with a constant rate of 5% CO2. At time points, supernatants were removed and neutrophils were washed with DPBS. Neutrophils were then incubated with 2% Saponin in PBS for 12 min and then dilution plated on THA to enumerate GBS CFU/well. To calculate %associated, the recovered CFU/well for all biological replicates of WT CJB111 at 4 hr were averaged and then each biological replicate for each strain was normalized to the WT average.

### Cytotoxicity assay

Cytotoxicity of HL60-neutrophils after infection with GBS strains was determined by LDH release using the CyQUANT LDH Cytotoxicity Assay (Invitrogen by Thermo Fisher C20301, Waltham, MA). Briefly, HL60-neutrophils were infected as described in the paragraph above and then at 5 hrs, 100 µL of supernatant from each well was removed and spun down at 500 x g for 1 min to remove cells and debris. Then the kit was performed per manufacture instructions.

### Flow cytometry detection of methylglyoxal-modified proteins in HL60 cells

To determine the impact of infection on methylglyoxal levels in HL60-neutrophils, differentiated Hl60s were first re-suspended in fresh RPMI + 10%FBS with 0 or 20mM glucose and allowed to equilibrate for 2 hrs. PBS or WT GBS at MOI 20 was added to 1 mL aliquots of 10^6^ differentiated HL60 cells and incubated at 37C with rotation for 2.5 hours and then harvested by centrifugation (500 x g). Cells were then stained using eBioscience Fixable Viability Dye eFluor 506 (Catalog # 65-0866-18) in PBS for 30 minutes at room temperature followed by anti-human Cd11b antibody conjugated to FITC (1:20 dilution; BD Biosciences 562793) in MACS buffer (1.25g BSA, 0.185g EDTA, 250mL PBS) for 30 minutes at room temperature. Cells were fixed and permeabilized using the FoxP3 fixation/permeabilization kit (Thermo Fisher Scientific, Catalog # 00-5523-00) according to manufacturer’s instructions before staining for intracellular methylglyoxal using an anti-MG antibody conjugated to PE (clone 9E7; Cat # MA5-45812; recognizes methylglyoxal-modified proteins) or an IgG2a isotype control (Cat # MG2A04) at final concentrations of 0.67 ug/mL (30 minutes in permeabilization buffer at room temperature). Stained cells were run on a BD LSRFortessa (BD Biosciences) using the BD FacsDiva software (v9) and analyzed by BD FlowJo software (v10.8). Gating strategy was determined by fluorescence minus one (FMO) controls. Flow cytometric histograms display anti-MG staining of 2000 CD11b+ events per sample.

### Statistical analysis

Statistical analysis was performed using Prism version 10.1 for Windows (GraphPad Software, San Diego, CA, USA) as described in figure legends.

## Acknowledgements

We would like to thank Dr. Jeffrey Kavanaugh from the University of Colorado Anschutz Medical Campus for help with protein modeling, and Dr. Sarah Stanley and Andrea Anaya Sanchez from the University of California Berkely for helpful discussions The work was supported by the American Heart Association grant 23POST1013835 to L.R.J. and the National Institutes of Health grants R01NS116716 and R01AI153332 to K.S.D., grant T32HD007186 to A.B. and grant F31AI178881 to M.S.A.

## Supplementary Material

**Table S1.** Complete *in vivo* blood Tn-sequencing dataset.

**Table S2.** Primers and Strains

**Figure S1**. The impact of MG on different GBS isolates. Growth curves for (A) CJB111 in mCDM with 0.5 mM and 1.0 mM MG quantified by CFU enumeration or (B) Growth curves for 10 GBS isolates in mCDM without MG (left) or with 1.0 mM MG (right) quantified by OD_600_.

**Figure S2**. GBS Glyoxalase A protein characterization. (A) Alignment of GloA amino acid sequences from GBS CJB111, GBS A909, GBS COH1, *S. pyogenes* 5448, and *E. coli* K-12. Green stars indicate known or predicted metal binding sites and colored bar indicates confidence of structure prediction. (B) Phylogenetic tree for 57 GBS GloA proteins. Proteins/branches with a mutation in amino acid residue A45 are labeled and colored. Red indicates an A45S mutation, purple indicates an A45T mutation, and grey indicates a mutation that only occurred once. (C) Superimposed tertiary structures for the solved *E. coli* GloA and the predicted GBS GloA. (D) idDT confidence scores across GBS GloA predicted structure. (E) Left: Predicted tertiary protein dimer for GBS A909/CJB111 GloA. Right: 180° horizontal rotation of the predicted tertiary protein dimer with the blue monomer containing GBS COH1 A45S mutation. Black arrows indicate predicted active site. (F) Baseline transcription of *gloA* and *gloB* genes grown to mid-log in mCDM and quantified by RT-qPCR. Gene transcript levels were normalized to the average CJB111 levels. Significance determined by 2way ANOVA with uncorrected Fisher’s LSD test, *P* < 0.05. * < 0.05, ** < 0.01.

**Figure S3.** The impact of MG and Glyoxalase A on GBS growth and virulence. (A) Growth curves for CJB111 in mCDM with or without MG after pre-exposure to 0.5 mM MG quantified by OD_600_. Growth curves for CJB111, Δ*gloA*, and p*gloA* strains in mCDM (B) with 10, 22, or 50 mM glucose quantified by OD_600_ or (C) with 0.1% (29.4mM) hydrogen peroxide quantified by CFU enumeration. (D) Percent red blood cell lysis of human blood for CJB111, Δ*gloA*, and p*gloA* strains relative to positive lysis control at 24hrs post-inoculation.

**Figure S4.** MG detoxification is more important as infection progresses. Recovered CFU counts from the blood of infected mice at (A) 6 and 24 hours post-infection and (B) TOD. Significance determined by (A) Mixed-effects analysis with uncorrected Fisher’s LSD test, or (B) Kruskal-Wallis with uncorrected Dunn’s test., *P* < 0.05. * < 0.05, ****<0.0001.

**Figure S5.** The impact of serum, primary cells, and glucose on neutrophil phenotypes. (A) Survival of CJB111, Δ*gloA*, and p*gloA* strains over time in the presence of serum without HL60-neutrophils. Percent survival was calculated by dividing CFU recovered from wells with GBS opsonized with normal serum by CFU recovered from wells with GBS opsonized with heat-killed (HK) serum. (B) Cytotoxicity of HL60-neutrophils incubated with CJB111, Δ*gloA*, and p*gloA* that had been opsonized with HK or normal serum after 5 hrs of infection. (C) Recovered CFU counts normalized to the WT average of WT CJB111, Δ*gloA*, and p*gloA* strains after 4 hrs of infection of BMNs. (D) Flow cytometry quantification of intracellular MG-modified proteins in HL60-neutrophils with or without WT CJB111 infection in 0mM glucose media. Left: Representative histogram displaying MG signal that includes isotype control, uninfected, and infected cells. Right: Geometric MFI quantification for MG in uninfected and infected samples. Significance determined by (C) one-way ANOVA with Uncorrected Fisher’s LSD test or (D) Unpaired Student *t* test, *P* < 0.05.

## Notes

https://www.ncbi.nlm.nih.gov/sra/?term=PRJNA1125445

### Competing Interest Statement

The authors have declared no competing interest.

### Summary of Updates

Major changes included characterization in murine bone marrow derived neutrophils, and complementing the gloA mutant in vivo. We added to or reformatted Figure 2, Figure 5, and Supplemental Figures 1-5.

## References

1. K. S. Doran, V. Nizet, Molecular pathogenesis of neonatal group B streptococcal infection: no longer in its infancy. Mol Microbiol 54, 23–31 (2004).

2. L. K. Francois Watkins et al., Epidemiology of Invasive Group B Streptococcal Infections Among Nonpregnant Adults in the United States, 2008-2016. JAMA Intern Med 179, 479–488 (2019).

3. A. A. El-Gendy et al., Serotyping and Antibiotic Susceptibility of Invasive Streptococcus agalactiae in Egyptian Patients with or without Diabetes Mellitus. Am J Trop Med Hyg 105, 1684–1689 (2021).

4. M. S. Edwards et al., Long-term sequelae of group B streptococcal meningitis in infants. J Pediatr 106, 717–722 (1985).

5. R. Libster et al., Long-term outcomes of group B streptococcal meningitis. Pediatrics 130, e8–15 (2012).

6. J. Hall et al., Maternal Disease With Group B Streptococcus and Serotype Distribution Worldwide: Systematic Review and Meta-analyses. Clin Infect Dis 65, S112–S124 (2017).

7. N. K. Kurian, D. Modi, Mechanisms of group B Streptococcus-mediated preterm birth: lessons learnt from animal models. Reprod Fertil 3, R109–R120 (2022).

8. A. Berardi et al., Understanding Factors in Group B Streptococcus Late-Onset Disease. Infect Drug Resist 14, 3207–3218 (2021).

9. E. M. Sabroske et al., Evolving antibiotic resistance in Group B Streptococci causing invasive infant disease: 1970-2021. Pediatr Res 93, 2067–2071 (2023).

10. J. Vornhagen, K. M. Adams Waldorf, L. Rajagopal, Perinatal Group B Streptococcal Infections: Virulence Factors, Immunity, and Prevention Strategies. Trends Microbiol 25, 919–931 (2017).

11. H. S. Manzer et al., The Group B Streptococcal Adhesin BspC Interacts with Host Cytokeratin 19 To Promote Colonization of the Female Reproductive Tract. mBio 13, e0178122 (2022).

12. B. L. Spencer et al., A type VII secretion system in Group B Streptococcus mediates cytotoxicity and virulence. PLoS Pathog 17, e1010121 (2021).

13. L. Mereghetti, I. Sitkiewicz, N. M. Green, J. M. Musser, Extensive adaptive changes occur in the transcriptome of Streptococcus agalactiae (group B streptococcus) in response to incubation with human blood. PLoS One 3, e3143 (2008).

14. M. S. Akbari, K. S. Doran, L. R. Burcham, Metal Homeostasis in Pathogenic Streptococci. Microorganisms 10, (2022).

15. Y. Liu, J. Liu, Group B Streptococcus: Virulence Factors and Pathogenic Mechanism. Microorganisms 10, (2022).

16. B. Spellerberg et al., Identification of genetic determinants for the hemolytic activity of Streptococcus agalactiae by ISS1 transposition. J Bacteriol 181, 3212–3219 (1999).

17. K. S. Doran, G. Y. Liu, V. Nizet, Group B streptococcal beta-hemolysin/cytolysin activates neutrophil signaling pathways in brain endothelium and contributes to development of meningitis. J Clin Invest 112, 736–744 (2003).

18. A. Ring et al., Group B streptococcal beta-hemolysin induces mortality and liver injury in experimental sepsis. J Infect Dis 185, 1745–1753 (2002).

19. G. Y. Liu et al., Sword and shield: linked group B streptococcal beta-hemolysin/cytolysin and carotenoid pigment function to subvert host phagocyte defense. Proc Natl Acad Sci U S A 101, 14491–14496 (2004).

20. M. E. Hensler et al., Virulence role of group B Streptococcus beta-hemolysin/cytolysin in a neonatal rabbit model of early-onset pulmonary infection. J Infect Dis 191, 1287–1291 (2005).

21. M. J. Cieslewicz et al., Structural and genetic diversity of group B streptococcus capsular polysaccharides. Infect Immun 73, 3096–3103 (2005).

22. K. Noble et al., Group B Streptococcus cpsE Is Required for Serotype V Capsule Production and Aids in Biofilm Formation and Ascending Infection of the Reproductive Tract during Pregnancy. ACS Infect Dis 7, 2686–2696 (2021).

23. A. Schuchat, Epidemiology of group B streptococcal disease in the United States: shifting paradigms. Clin Microbiol Rev 11, 497–513 (1998).

24. L. Madrid et al., Infant Group B Streptococcal Disease Incidence and Serotypes Worldwide: Systematic Review and Meta-analyses. Clin Infect Dis 65, S160–S172 (2017).

25. H. J. Jennings, C. Lugowski, D. L. Kasper, Conformational aspects critical to the immunospecificity of the type III group B streptococcal polysaccharide. Biochemistry 20, 4511–4518 (1981).

26. M. S. Edwards, D. L. Kasper, H. J. Jennings, C. J. Baker, A. Nicholson-Weller, Capsular sialic acid prevents activation of the alternative complement pathway by type III, group B streptococci. J Immunol 128, 1278–1283 (1982).

27. J. Hayrinen, S. Pelkonen, J. Finne, Structural similarity of the type-specific group B streptococcal polysaccharides and the carbohydrate units of tissue glycoproteins: evaluation of possible cross-reactivity. Vaccine 7, 217–224 (1989).

28. J. Morgenstern, M. Campos Campos, P. Nawroth, T. Fleming, The Glyoxalase System-New Insights into an Ancient Metabolism. Antioxidants (Basel) 9, (2020).

29. M. J. MacLean, L. S. Ness, G. P. Ferguson, I. R. Booth, The role of glyoxalase I in the detoxification of methylglyoxal and in the activation of the KefB K+ efflux system in Escherichia coli. Mol Microbiol 27, 563–571 (1998).

30. S. W. T. Lai, E. J. Lopez Gonzalez, T. Zoukari, P. Ki, S. C. Shuck, Methylglyoxal and Its Adducts: Induction, Repair, and Association with Disease. Chem Res Toxicol 35, 1720–1746 (2022).

31. B. L. Spencer, A. Chatterjee, B. A. Duerkop, C. J. Baker, K. S. Doran, Complete Genome Sequence of Neonatal Clinical Group B Streptococcal Isolate CJB111. Microbiol Resour Announc 10, (2021).

32. L. R. Burcham et al., Genomic Analyses Identify Manganese Homeostasis as a Driver of Group B Streptococcal Vaginal Colonization. mBio 13, e0098522 (2022).

33. L. R. Burcham et al., Identification of Zinc-Dependent Mechanisms Used by Group B Streptococcus To Overcome Calprotectin-Mediated Stress. mBio 11, (2020).

34. Y. Inoue, A. Kimura, Methylglyoxal and regulation of its metabolism in microorganisms. Adv Microb Physiol 37, 177–227 (1995).

35. C. Poyart et al., Attenuated virulence of Streptococcus agalactiae deficient in D-alanyl-lipoteichoic acid is due to an increased susceptibility to defensins and phagocytic cells. Mol Microbiol 49, 1615–1625 (2003).

36. Q. Cheng, D. Stafslien, S. S. Purushothaman, P. Cleary, The group B streptococcal C5a peptidase is both a specific protease and an invasin. Infect Immun 70, 2408–2413 (2002).

37. S. Dramsi et al., Assembly and role of pili in group B streptococci. Mol Microbiol 60, 1401–1413 (2006).

38. K. S. Doran et al., Blood-brain barrier invasion by group B Streptococcus depends upon proper cell-surface anchoring of lipoteichoic acid. J Clin Invest 115, 2499–2507 (2005).

39. C. Whidbey et al., A hemolytic pigment of Group B Streptococcus allows bacterial penetration of human placenta. J Exp Med 210, 1265–1281 (2013).

40. E. Boldenow et al., Group B Streptococcus circumvents neutrophils and neutrophil extracellular traps during amniotic cavity invasion and preterm labor. Sci Immunol 1, (2016).

41. S. Papasergi et al., The GBS PI-2a pilus is required for virulence in mice neonates. PLoS One 6, e18747 (2011).

42. K. A. Patras et al., Group B Streptococcus Biofilm Regulatory Protein A Contributes to Bacterial Physiology and Innate Immune Resistance. J Infect Dis 218, 1641–1652 (2018).

43. V. Mercado-Evans et al., Gestational diabetes augments group B Streptococcus infection by disrupting maternal immunity and the vaginal microbiota. Nat Commun 15, 1035 (2024).

44. A. F. Carlin et al., Group B Streptococcus suppression of phagocyte functions by protein-mediated engagement of human Siglec-5. J Exp Med 206, 1691–1699 (2009).

45. S. Uchiyama et al., Dual actions of group B Streptococcus capsular sialic acid provide resistance to platelet-mediated antimicrobial killing. Proc Natl Acad Sci U S A 116, 7465–7470 (2019).

46. P. Moulin et al., The Adc/Lmb System Mediates Zinc Acquisition in Streptococcus agalactiae and Contributes to Bacterial Growth and Survival. J Bacteriol 198, 3265–3277 (2016).

47. M. S. Akbari et al., The impact of nutritional immunity on Group B streptococcal pathogenesis during wound infection. mBio, e0030423 (2023).

48. A. Clancy et al., Evidence for siderophore-dependent iron acquisition in group B streptococcus. Mol Microbiol 59, 707–721 (2006).

49. S. Shabayek, R. Bauer, S. Mauerer, B. Mizaikoff, B. Spellerberg, A streptococcal NRAMP homologue is crucial for the survival of Streptococcus agalactiae under low pH conditions. Mol Microbiol 100, 589–606 (2016).

50. M. J. Sullivan, K. G. K. Goh, D. Gosling, L. Katupitiya, G. C. Ulett, Copper Intoxication in Group B Streptococcus Triggers Transcriptional Activation of the cop Operon That Contributes to Enhanced Virulence during Acute Infection. J Bacteriol 203, e0031521 (2021).

51. C. Faralla et al., Analysis of two-component systems in group B Streptococcus shows that RgfAC and the novel FspSR modulate virulence and bacterial fitness. mBio 5, e00870–00814 (2014).

52. L. Thomas, L. Cook, Two-Component Signal Transduction Systems in the Human Pathogen Streptococcus agalactiae. Infect Immun 88, (2020).

53. D. S. Ipe et al., Discovery and Characterization of Human-Urine Utilization by Asymptomatic-Bacteriuria-Causing Streptococcus agalactiae. Infect Immun 84, 307–319 (2016).

54. C. Poyart, M. C. Lamy, C. Boumaila, F. Fiedler, P. Trieu-Cuot, Regulation of D-alanyl-lipoteichoic acid biosynthesis in Streptococcus agalactiae involves a novel two-component regulatory system. J Bacteriol 183, 6324–6334 (2001).

55. D. Quach et al., The CiaR response regulator in group B Streptococcus promotes intracellular survival and resistance to innate immune defenses. J Bacteriol 191, 2023–2032 (2009).

56. Y. Yang et al., Role of Two-Component System Response Regulator bceR in the Antimicrobial Resistance, Virulence, Biofilm Formation, and Stress Response of Group B Streptococcus. Front Microbiol 10, 10 (2019).

57. S. Khosa, Z. AlKhatib, S. H. Smits, NSR from Streptococcus agalactiae confers resistance against nisin and is encoded by a conserved nsr operon. Biol Chem 394, 1543–1549 (2013).

58. S. Khosa et al., Structural basis of lantibiotic recognition by the nisin resistance protein from Streptococcus agalactiae. Sci Rep 6, 18679 (2016).

59. L. Rajagopal, A. Vo, A. Silvestroni, C. E. Rubens, Regulation of purine biosynthesis by a eukaryotic-type kinase in Streptococcus agalactiae. Mol Microbiol 56, 1329–1346 (2005).

60. G. S. Tamura, A. Nittayajarn, D. L. Schoentag, A glutamine transport gene, glnQ, is required for fibronectin adherence and virulence of group B streptococci. Infect Immun 70, 2877–2885 (2002).

61. D. S. Ipe et al., Conserved bacterial de novo guanine biosynthesis pathway enables microbial survival and colonization in the environmental niche of the urinary tract. ISME J 15, 2158–2162 (2021).

62. I. Santi et al., CsrRS regulates group B Streptococcus virulence gene expression in response to environmental pH: a new perspective on vaccine development. J Bacteriol 191, 5387–5397 (2009).

63. Q. Yang, M. Zhang, D. J. Harrington, G. W. Black, I. C. Sutcliffe, A proteomic investigation of Streptococcus agalactiae reveals that human serum induces the C protein beta antigen and arginine deiminase. Microbes Infect 13, 757–760 (2011).

64. I. van de Rijn, R. E. Kessler, Growth characteristics of group A streptococci in a new chemically defined medium. Infect Immun 27, 444–448 (1980).

65. M. M. Zhang, C. L. Ong, M. J. Walker, A. G. McEwan, Defence against methylglyoxal in Group A Streptococcus: a role for Glyoxylase I in bacterial virulence and survival in neutrophils? Pathog Dis 74, (2016).

66. A. Anaya-Sanchez, Y. Feng, J. C. Berude, D. A. Portnoy, Detoxification of methylglyoxal by the glyoxalase system is required for glutathione availability and virulence activation in Listeria monocytogenes. PLoS Pathog 17, e1009819 (2021).

67. M. M. He, S. L. Clugston, J. F. Honek, B. W. Matthews, Determination of the structure of Escherichia coli glyoxalase I suggests a structural basis for differential metal activation. Biochemistry 39, 8719–8727 (2000).

68. L. A. O’Neill, R. J. Kishton, J. Rathmell, A guide to immunometabolism for immunologists. Nat Rev Immunol 16, 553–565 (2016).

69. K. H. Darwin, S. A. Stanley, The aldehyde hypothesis: metabolic intermediates as antimicrobial effectors. Open Biol 12, 220010 (2022).

70. G. Limon, N. M. Samhadaneh, A. Pironti, K. H. Darwin, Aldehyde accumulation in Mycobacterium tuberculosis with defective proteasomal degradation results in copper sensitivity. mBio 14, e0036323 (2023).

71. H. Rachman et al., Critical role of methylglyoxal and AGE in mycobacteria-induced macrophage apoptosis and activation. PLoS One 1, e29 (2006).

72. J. H. Jeon, C. W. Hong, E. Y. Kim, J. M. Lee, Current Understanding on the Metabolism of Neutrophils. Immune Netw 20, e46 (2020).

73. J. M. Daley, A. A. Thomay, M. D. Connolly, J. S. Reichner, J. E. Albina, Use of Ly6G-specific monoclonal antibody to deplete neutrophils in mice. J Leukoc Biol 83, 64–70 (2008).

74. R. L. Goldenberg, J. C. Hauth, W. W. Andrews, Intrauterine infection and preterm delivery. N Engl J Med 342, 1500–1507 (2000).

75. T. A. Hooven et al., The Streptococcus agalactiae Stringent Response Enhances Virulence and Persistence in Human Blood. Infect Immun 86, (2018).

76. L. Mereghetti, I. Sitkiewicz, N. M. Green, J. M. Musser, Identification of an unusual pattern of global gene expression in group B Streptococcus grown in human blood. PLoS One 4, e7145 (2009).

77. B. Di Palo et al., Adaptive response of Group B streptococcus to high glucose conditions: new insights on the CovRS regulation network. PLoS One 8, e61294 (2013).

78. Y. Lopez et al., Serotype, virulence profile, antimicrobial resistance and macrolide-resistance determinants in Streptococcus agalactiae isolates in pregnant women and neonates in Catalonia, Spain. Enferm Infecc Microbiol Clin (Engl Ed) 36, 472–477 (2018).

79. X. Zhang, C. G. Schalkwijk, K. Wouters, Immunometabolism and the modulation of immune responses and host defense: A role for methylglyoxal? Biochim Biophys Acta Mol Basis Dis 1868, 166425 (2022).

80. P. J. Beisswenger, Methylglyoxal in diabetes: link to treatment, glycaemic control and biomarkers of complications. Biochem Soc Trans 42, 450–456 (2014).

81. C. Lee, C. Park, Bacterial Responses to Glyoxal and Methylglyoxal: Reactive Electrophilic Species. Int J Mol Sci 18, (2017).

82. S. L. Hazen, F. F. Hsu, A. d’Avignon, J. W. Heinecke, Human Neutrophils Employ Myeloperoxidase To Convert a-Amino Acids to a Batter of Reactive Aldehydes: A Pathway for Aldehyde Generation at Sites of Inflammation. Biochemistry 37, 6864–6873 (1998).

83. A. Furuta et al., Bacterial and Host Determinants of Group B Streptococcal Infection of the Neonate and Infant. Front Microbiol 13, 820365 (2022).

84. S. L. Clugston, R. Yajima, J. F. Honek, Investigation of metal binding and activation of Escherichia coli glyoxalase I: kinetic, thermodynamic and mutagenesis studies. Biochem J 377, 309–316 (2004).

85. U. Suttisansanee et al., Structural variation in bacterial glyoxalase I enzymes: investigation of the metalloenzyme glyoxalase I from Clostridium acetobutylicum. J Biol Chem 286, 38367–38374 (2011).

86. L. Zeng, P. Noeparvar, R. A. Burne, B. S. Glezer, Genetic characterization of glyoxalase pathway in oral streptococci and its contribution to interbacterial competition. J Oral Microbiol 16, 2322241 (2024).

87. L. C. C. Cook, H. Hu, M. Maienschein-Cline, M. J. Federle, A Vaginal Tract Signal Detected by the Group B Streptococcus SaeRS System Elicits Transcriptomic Changes and Enhances Murine Colonization. Infect Immun 86, (2018).

88. M. Alfano, C. Cavazza, Structure, function, and biosynthesis of nickel-dependent enzymes. Protein Sci 29, 1071–1089 (2020).

89. T. Eitinger, M. A. Mandrand-Berthelot, Nickel transport systems in microorganisms. Arch Microbiol 173, 1–9 (2000).

90. N. Krymkiewicz, E. Dieguez, U. D. Rekarte, N. Zwaig, Properties and mode of action of a bactericidal compound (=methylglyoxal) produced by a mutant of Escherichia coli. J Bacteriol 108, 1338–1347 (1971).

91. K. Takahashi, Further studies on the reactions of phenylglyoxal and related reagents with proteins. J Biochem 81, 403–414 (1977).

92. K. Takahashi, The reactions of phenylglyoxal and related reagents with amino acids. J Biochem 81, 395–402 (1977).

93. S. T. Cheung, M. L. Fonda, Kinetics of the inactivation of Escherichia coli glutamate apodecarboxylase by phenylglyoxal. Arch Biochem Biophys 198, 541–547 (1979).

94. N. Rabbani, P. J. Thornalley, Measurement of methylglyoxal by stable isotopic dilution analysis LC-MS/MS with corroborative prediction in physiological samples. Nat Protoc 9, 1969–1979 (2014).

95. C. G. Schalkwijk, C. D. A. Stehouwer, Methylglyoxal, a Highly Reactive Dicarbonyl Compound, in Diabetes, Its Vascular Complications, and Other Age-Related Diseases. Physiol Rev 100, 407–461 (2020).

96. R. A. Keogh et al., Group B Streptococcus adaptation promotes survival in a hyperinflammatory diabetic wound environment. Sci Adv 8, eadd3221 (2022).

97. S. Kant, L. Liu, A. Vazquez-Torres, The methylglyoxal pathway is a sink for glutathione in Salmonella experiencing oxidative stress. PLoS Pathog 19, e1011441 (2023).

98. L. R. Joyce et al., Identification of a novel cationic glycolipid in Streptococcus agalactiae that contributes to brain entry and meningitis. PLoS Biol 20, e3001555 (2022).

99. A. Banerjee et al., Bacterial Pili exploit integrin machinery to promote immune activation and efficient blood-brain barrier penetration. Nat Commun 2, 462 (2011).

100. B. J. Kim et al., Bacterial induction of Snail1 contributes to blood-brain barrier disruption. J Clin Invest 125, 2473–2483 (2015).

101. B. L. Spencer et al., Cas9 Contributes to Group B Streptococcal Colonization and Disease. Front Microbiol 10, 1930 (2019).

102. S. W. Wingett, S. Andrews, FastQ Screen: A tool for multi-genome mapping and quality control. F1000Res 7, 1338 (2018).

103. M. Martin, Cutadapt removes adapter sequences from high-throughput sequencing reads. 2011 (10.14806/ej.17.1.200).

104. M. A. DeJesus, C. Ambadipudi, R. Baker, C. Sassetti, T. R. Ioerger, TRANSIT--A Software Tool for Himar1 TnSeq Analysis. PLoS Comput Biol 11, e1004401 (2015).

105. H. Li, R. Durbin, Fast and accurate short read alignment with Burrows-Wheeler transform. Bioinformatics 25, 1754–1760 (2009).

106. F. Madeira et al., Search and sequence analysis tools services from EMBL-EBI in 2022. Nucleic Acids Res 50, W276–W279 (2022).

107. X. Robert, P. Gouet, Deciphering key features in protein structures with the new ENDscript server. Nucleic Acids Res 42, W320–324 (2014).

108. S. F. Altschul et al., Gapped BLAST and PSI-BLAST: a new generation of protein database search programs. Nucleic Acids Res 25, 3389–3402 (1997).

109. S. F. Altschul et al., Protein database searches using compositionally adjusted substitution matrices. FEBS J 272, 5101–5109 (2005).

110. J. Jumper et al., Highly accurate protein structure prediction with AlphaFold. Nature 596, 583–589 (2021).

111. M. Mirdita et al., ColabFold: making protein folding accessible to all. Nat Methods 19, 679–682 (2022).

112. H. M. Berman et al., The Protein Data Bank. Nucleic Acids Res 28, 235–242 (2000).

113. L. R. Joyce et al., Streptococcus agalactiae glycolipids promote virulence by thwarting immune cell clearance. Sci Adv 10, eadn7848 (2024).

