## Supplementary figures and images for "Identification of Glyoxalase A in Group B *Streptococcus* and its contribution to methylglyoxal tolerance and virulence"

### Supplemental Figure 1

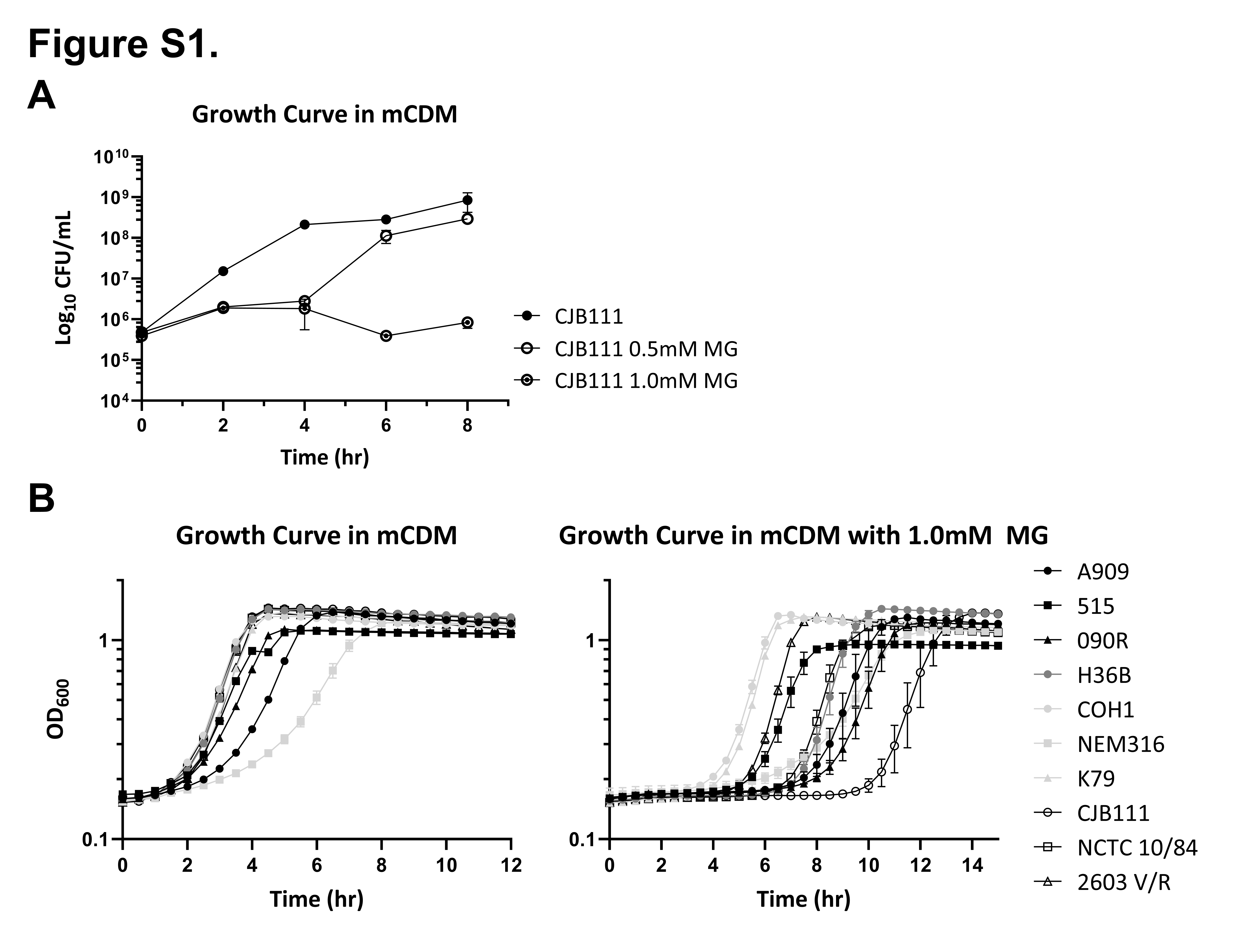

### Supplemental Figure 2

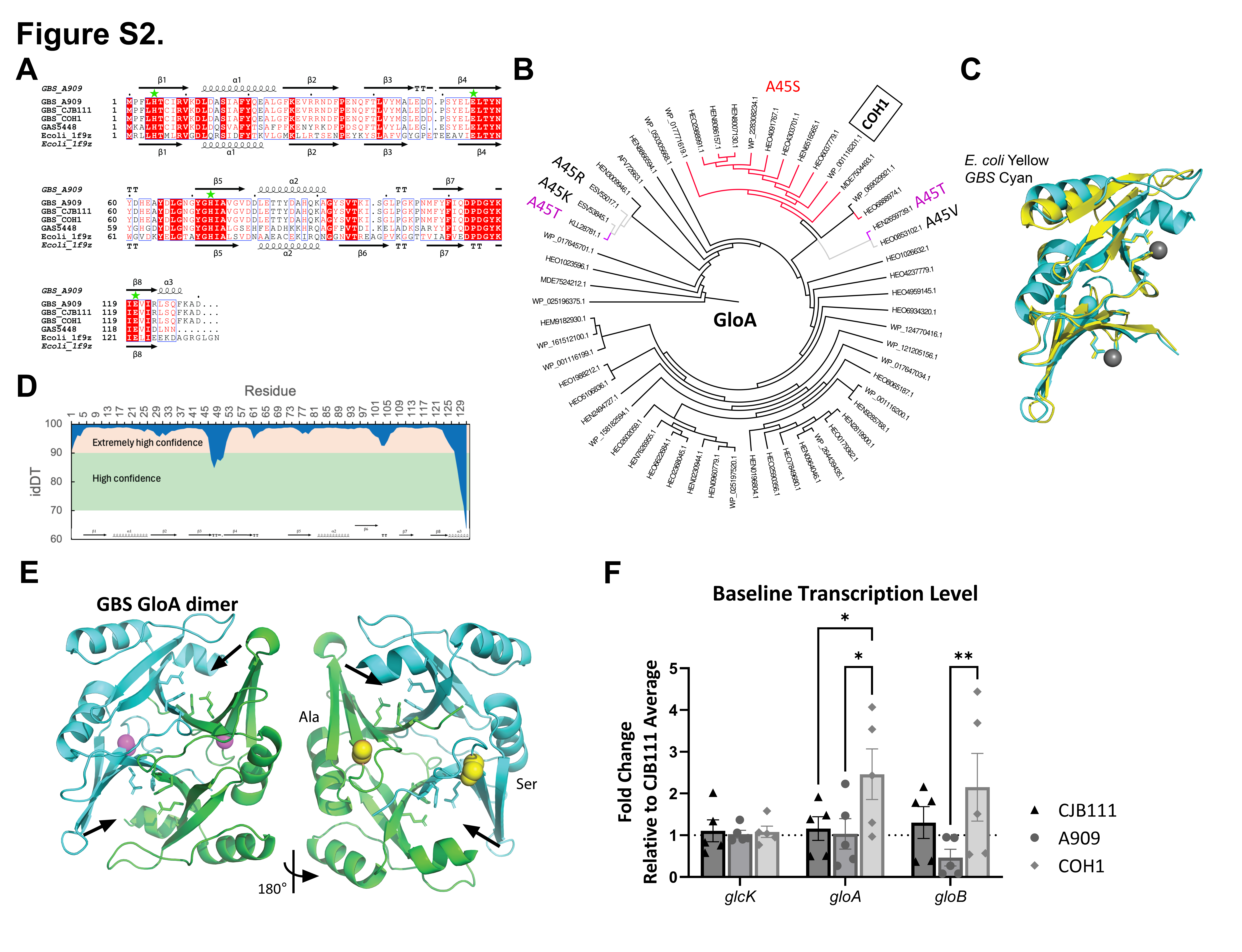

### Supplemental Figure 3

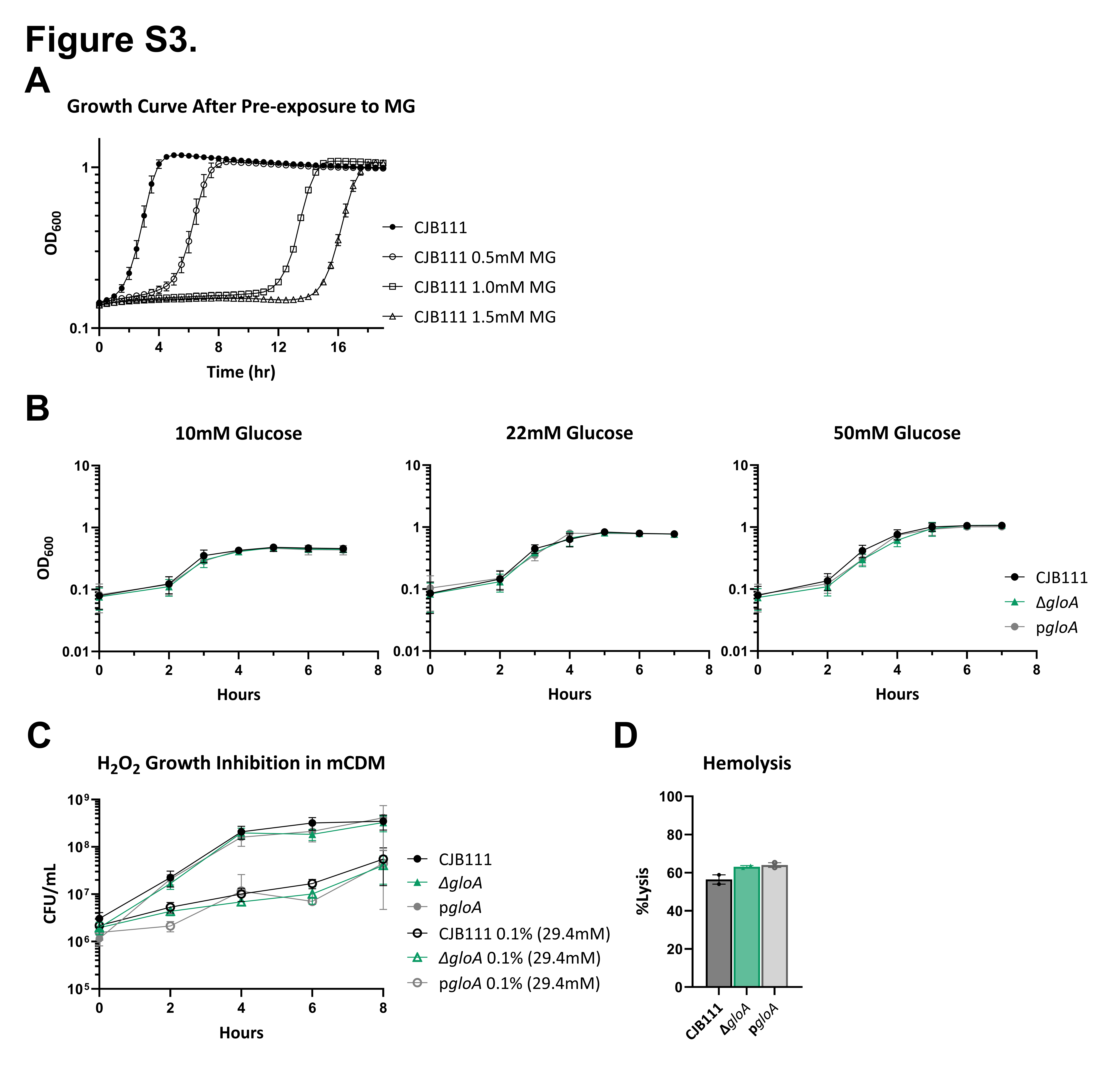

### Supplemental Figure 4

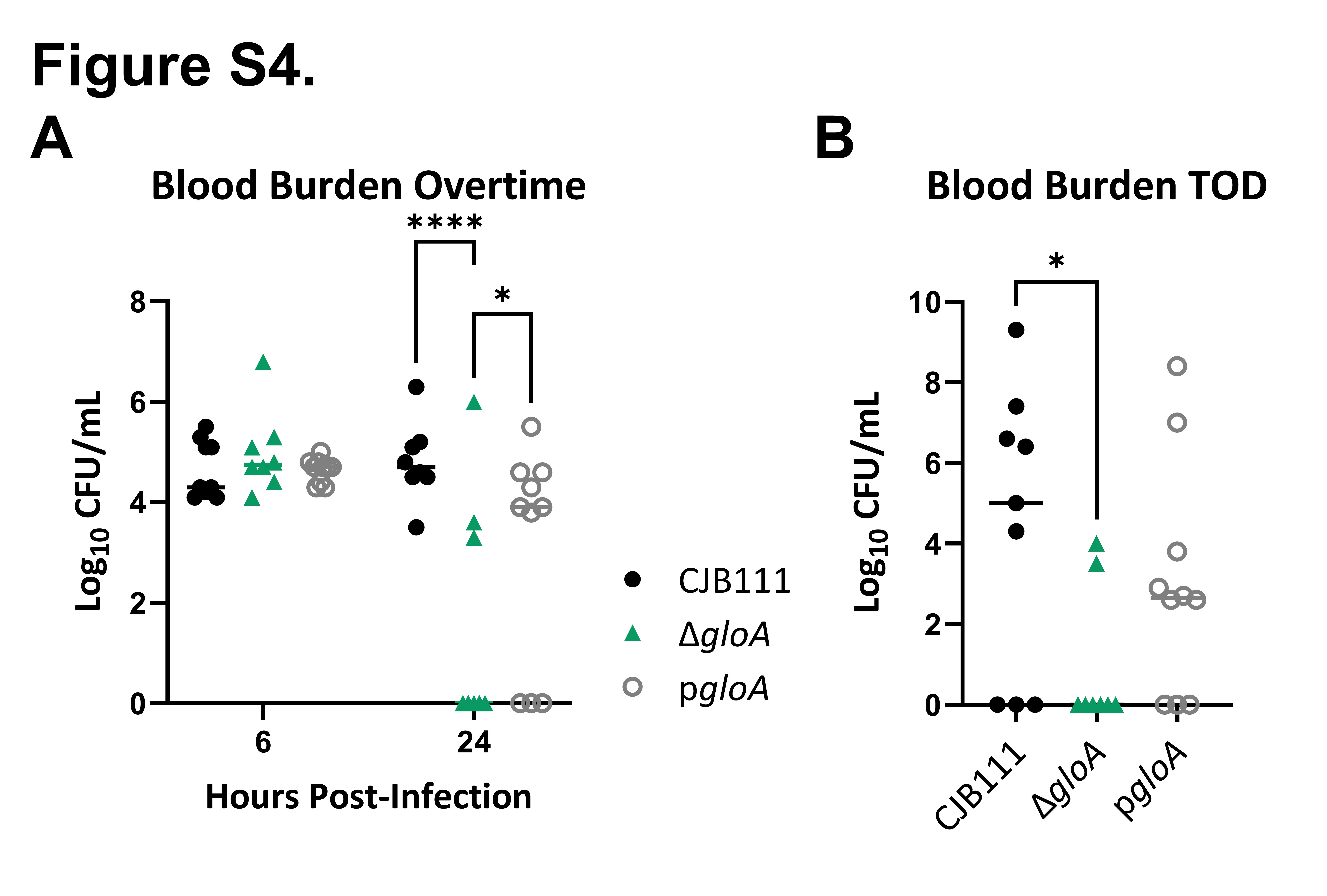

### Supplemental Figure 5

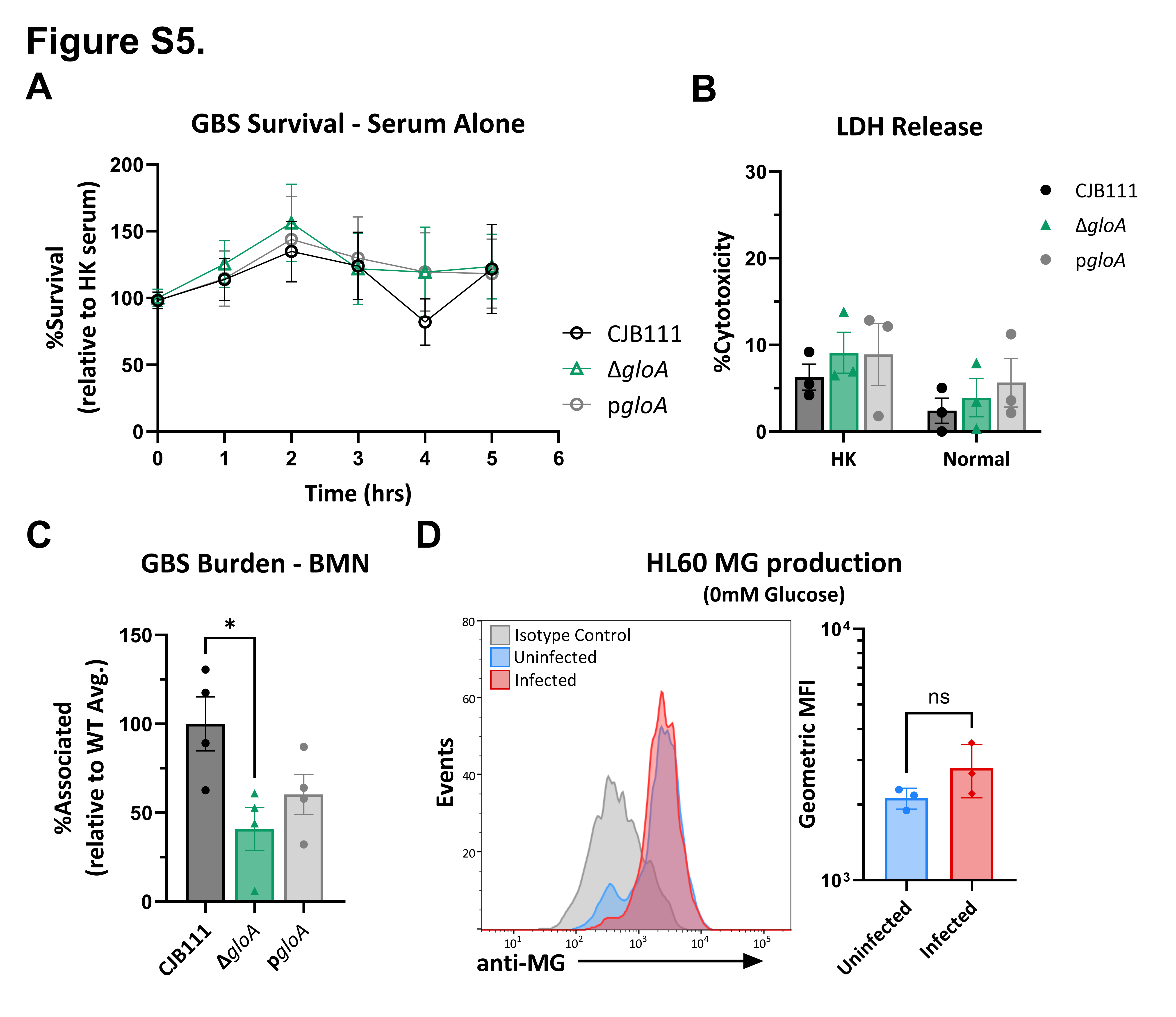
